# Engineered 3D Immuno-Glial-Neurovascular Human miBrain Model

**DOI:** 10.1101/2023.08.15.553453

**Authors:** Alice E. Stanton, Adele Bubnys, Emre Agbas, Benjamin James, Dong Shin Park, Alan Jiang, Rebecca L. Pinals, Liwang Liu, Nhat Truong, Anjanet Loon, Colin Staab, Oyku Cerit, Hsin-Lan Wen, Manolis Kellis, Joel W. Blanchard, Robert Langer, Li-Huei Tsai

**Author notes:** Department of Neuroscience, Black Family Stem Cell Institute, Ronald M. Loeb Center for Alzheimer’s Disease, Icahn School of Medicine at Mt. Sinai, New York, NY 10029. These authors contributed equally to this work (Alice E. Stanton, Adele Bubnys).

## Abstract

Patient-specific, human-based cellular models integrating a biomimetic blood-brain barrier (BBB), immune, and myelinated neuron components are critically needed to enable accelerated, translationally relevant discovery of neurological disease mechanisms and interventions. By engineering a novel brain-mimicking 3D hydrogel and co-culturing all six major brain cell types derived from patient iPSCs, we have constructed, characterized, and utilized a multicellular integrated brain (miBrain) immuno-glial-neurovascular model with *in vivo-*like hallmarks inclusive of neuronal activity, functional connectivity, barrier function, myelin-producing oligodendrocyte engagement with neurons, multicellular interactions, and transcriptomic profiles. We implemented the model to study Alzheimer’s Disease pathologies associated with *APOE4* genetic risk. *APOE4* miBrains differentially exhibit amyloid aggregation, tau phosphorylation, and astrocytic GFAP. Unlike the co-emergent fate specification of glia and neurons in organoids, miBrains integrate independently differentiated cell types, a feature we harnessed to identify that *APOE4* in astrocytes promotes neuronal tau pathogenesis and dysregulation through crosstalk with microglia.

## Introduction

*In vitro* brain models hold enormous potential for decoding disease mechanisms and establishing enhanced drug screening platforms for the discovery and development of therapeutic interventions^1–6^. Neurological disease treatments have been particularly hindered by the lack of human-based models, as genetic differences limit the pace and fidelity of therapeutic translation from rodent models to human^7^, many genetic risk variants lie in non-coding regions, and there are marked differences between the human brain and that of other species^8^. Induced pluripotent stem cell (iPSC) technology enables the generation of human patient-specific cells into almost any cell type in the body^9,10^. More recently, brain organoids, harnessing iPSCs, have enhanced neuronal maturation beyond that attainable in neuronal monoculture and provide a model of neuronal development with an emergent approach whereby neuronal progenitors differentiate together into a dense co-culture of neurons and glia with a maintained progenitor population with diverse signatures^1–3,11^. The utility of brain organoids has been further broadened by approaches to integrate microglial components^12,13^ by some groups, and vascular-like cells^14,15^ by others, though stable integration of all of these components, along with myelinating oligodendroglia has yet to be demonstrated. The dense cellular organization, which also often leads to the formation of a necrotic core, further limits the homogeneous distribution and biomimetic integration of cellular components. At the same time, others have taken more directed approaches to construct microphysiological systems with tissue-mimicking 3D architecture in which cells of the desired lineages are combined in a 3D scaffold via cell self-assembly or via patterning^4,5,16–20^.

Given the key role of glial, immune, and vascular cells in healthy brain physiological processes and that non-cell autonomous effects contribute critically to neurodegenerative disease^21–25^, there is great interest in establishing vascularized human brain models with all relevant CNS cell types and biomimetic tissue architecture. Further, to identify cell type- and pathway-specific contributions to pathogenesis, a brain model in which each cell type could be independently modified is desirable. To realize these possibilities, we have developed a 3D, iPSC-based multicellular integrated brain (miBrain) immuno-glial-neurovascular model. As conventional 3D microphysiological systems utilize protein-based materials that mimic only the basement membrane (BM), such as Matrigel and collagen, or rely on fibrin, like which forms during blood clots and contains neurotoxic components^26^, all of which rapidly degrade, we critically need a hydrogel mimetic of brain tissue beyond the BM, one which can also promote neuronal activity, while providing 3D structure and supporting cellular self-assembly. To this end, we engineered a novel 3D hydrogel, Neuromatrix Hydrogel, to provide enhanced brain mimicry and promote co-self-assembly of all six major CNS cell types into an integral engineered tissue. As one proof of principle, we have leveraged the miBrain to investigate the strongest genetic risk factor for sporadic Alzheimer’s Disease (AD), apolipoprotein E4 *(APOE4)*^27^. Importantly, we have modeled amyloid and tau pathologies in this integrated brain model and here uncovered that *APOE4* astrocytes are sufficient to induce reactive species and tau phosphorylation via crosstalk with microglia.

## Results

### Multicellular Integrated miBrain Model Co-Assembling 6 Major CNS Cell Types

Many human neurological disease involve a diversity of cell types beyond neurons, but there are currently no models that have both this full cellular diversity and human genetics^28^. We endeavored to address this critical void by developing a platform that is inclusive of all six CNS cell types, recapitulates important features of 3D brain tissue structure, and decouples glial and neuronal fate specification, enabling introduction of cell type-specific perturbations and genetic mutations. To this end, we separately differentiated, optimized, and validated each of the six major brain cell types from patient-specific iPSCs for: brain microvascular endothelial cells (BMECs)^29^, pericytes^29,30^, oligodendrocyte precursor cells (OPCs)^31^, neurons^32^, astrocytes^33^, and microglia-like cells (iMG)^34^.

We validated that iPSC-derived BMECs express canonical markers via immunohistochemistry (**Fig. S1A,B**), gene expression (**Fig. S1C,D**), and flow cytometry (**Fig. S2Ai**). As evidence suggests that some endothelial cell differentiation protocols from iPSCs can lead to more epithelial-like lineages^35^, we performed RNA-sequencing (RNAseq) on the BMECs used in this study and integrated the data with existing literature datasets^35–40^. We found that our BMECs clustered with endothelial cells—upregulated endothelial processes (**Fig. S1Ci**) and downregulated epithelial processes (**Fig. S1Cii**)—with comparable or greater expression of endothelial genes (**Fig. S1Di**) and lesser gene expression of epithelial genes (**Fig. S1Dii**) across the corresponding key gene sets^35^. Functionally, barrier resistance values for these BMECs were comparable to reported values for endothelial cells, as opposed to the much higher resistance in epithelial monolayers^35^ (**Fig. S1E**).

iPSC-derived pericytes were validated for protein expression of CD140b via flow cytometry (**Fig. S2Aii**), NG2 and PDGFRβ via immunoreactivity (**Fig. S2B,C**), as well as for gene expression for canonical marker genes, such as *PDGFRβ, CSPG4,* and *VEGFA* (**Fig. S3Ai**) and genes shown to be upregulated in pericytes and downregulated in smooth muscle cells^41^, such as *TGFβ1, DCHS1,* and *NUAK1* (**Fig. S3Aii**). iPSC-derived astrocytes displayed strong protein expression of CD44 via flow cytometry (**Fig. S2Aiii**), S100β, GFAP, ALDH1L1, and AQP4 via immunoreactivity (**Fig. S2B,D**), a lack of immunoreactivity to neuron progenitor marker SOX2 (**Fig. S2B**), positive gene expression for canonical markers like *VIM* and *GFAP* (**Fig. S3B**), and, functionally, calcium transients (**Fig. S4**). iPSC-derived OPCs demonstrated PDGFRα protein expression via flow cytometry (**Fig. S2Aiv**), SOX10, Olig2, O4, and PDFRα immunoreactivity (**Fig. S2B,E**), and expression for identity genes *like PAX6, PDGFRα*, *and PLP1* (**Fig. S3C**). iMG were validated for CD45 marker expression via flow cytometry (**Fig. S2Avi**), immunoreactivity to Iba1, P2RY12, and TMEM119 (**Fig. S2B,G**), and expression for identity genes like *P2RY12, FOS,* and *CSF1R* (**Fig. S3E**). iPSC-derived neurons displayed strong expression of excitatory cortical neuron marker PSA NCAM via flow cytometry (**Fig. S2Av**), β-Tubulin, SOX2, synapsin, MAP2, and NeuN immunoreactivity, and a lack of immunoreactivity to pluripotency marker Oct4, here included because NGN2-neurons rely on piggyBac-mediated expression of the cassette, without which pluripotent cells could persistent (**Fig. S2B,F**), and canonical genes such as *MAP2, SCN1A,* and *NTRK2* (**Fig. S3D**).

Upon validated fate specification for each lineage, cells were combined into a novel 3D hydrogel we engineered to mimic key aspects of brain tissue, enabling 3D cell co-culture and network co-assembly, and seeded in 48-well plates for high throughput formation of the engineered miBrains (**Fig. 1A**). miBrains are characterized by their composition, including all six major CNS cell types together, distributed throughout the 3D tissue, and their emergent 3D morphologies and functionalities.

**Fig. 1:**
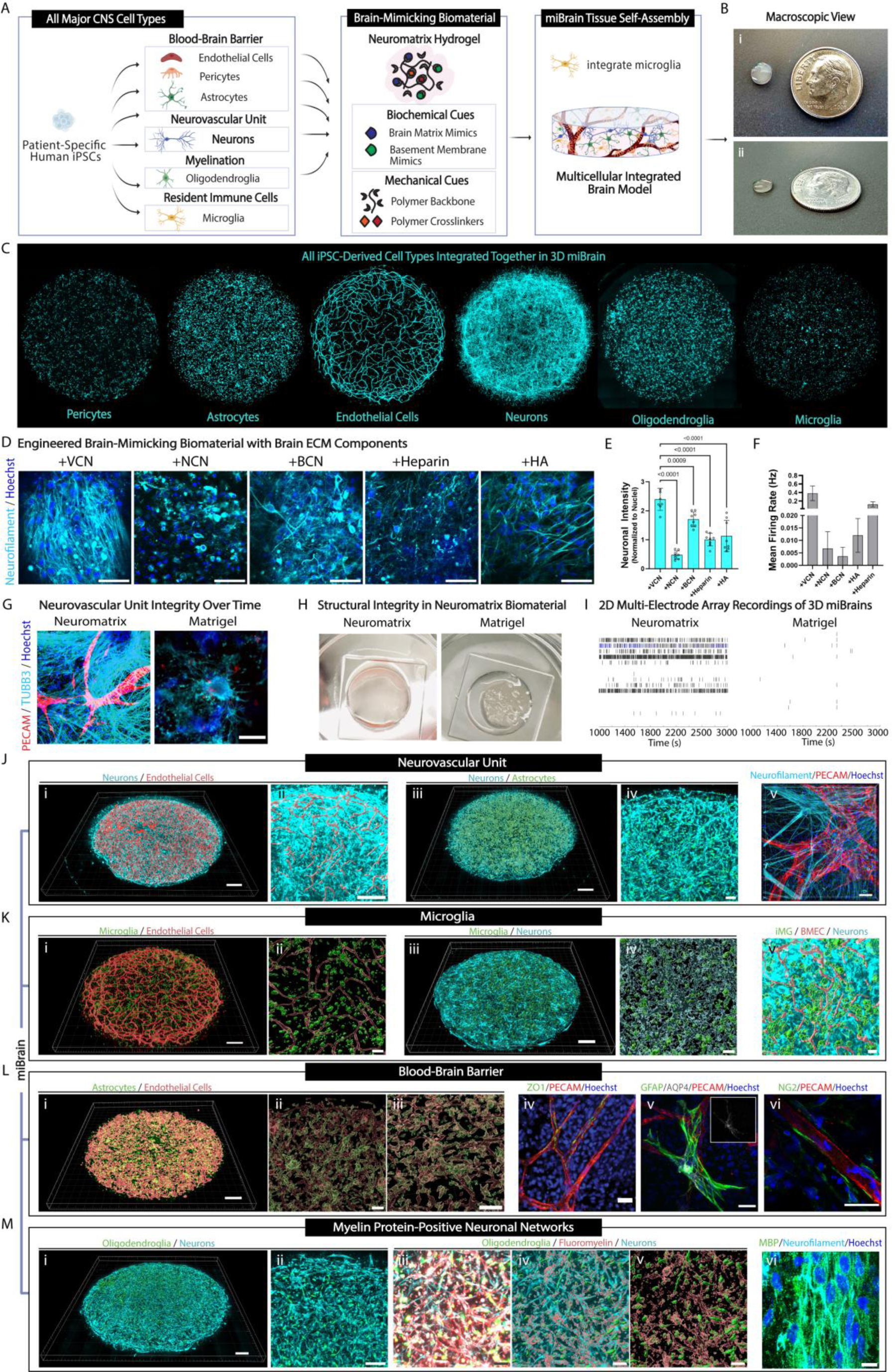
Human Integrated 3D-Immuno-Glial-Neurovascular miBrain Model. (**A**) Schematic of miBrain formation harnessing patient-specific iPSCs differentiated into each of the resident brain cell types, encapsulated in Neuromatrix Hydrogel, and co-cultured for integral cell network self-assembly and microglia-like cell integration, (**B**) macroscopic view of miBrains from (i) top and (ii) side angle pictured with a dime for reference, (**C**) distribution of (from left to right) iPSC-derived pericytes (cyan: mCherry-pericytes), astrocytes (cyan: mCherry-astrocytes), BMECs (cyan: mCherry-BMECs), neurons (cyan: tubulin neuron label), oligodendroglia (cyan: tdTomato pre-transfected oligodendroglia), and iMG (cyan: membrane pre-labeled iMG) throughout the full 3D miBrain, (**D**) neuronal phenotypes in miBrains cultured in dextran-based hydrogels fabricated with various brain ECM proteins (cyan: neurofilament, blue: Hoechst; scale bars, 50 µm), (**E**) quantification of neurofilament immunoreactivity (averages of n = 8 samples from n = 3 fields of view per sample), (**F**) neuronal firing as assessed on an MEA system across conditions (n = 3 wells per group, displayed as mean and S.E.M.), (**G**) persistence of neurovascular unit phenotypes in miBrains cultured in Neuromatrix Hydrogel versus Matrigel after 5 weeks in culture (red: PECAM, cyan: TUBB3, blue: Hoechst; scale bar, 50 µm), (**H**) macroscopic view of gel structural integrity for miBrains cultured in VCN-incorporated engineered dextran-based hydrogel named Neuromatrix Hydrogel versus Matrigel after 4 weeks, (**I**) example raster plots from MEA recordings of miBrains in Neuromatrix Hydrogel compared to Matrigel; miBrains recapitulate key hallmarks of human brain tissue, inclusive of (**J**) neurovascular units: (i) 3D integrated BMEC and neuronal networks throughout the miBrain (red: Imaris reconstruction of mCherry-BMECs, cyan: tubulin neuron label, scale bar, 500 µm), (ii) visualized at higher magnification (red: Imaris surfaces of mCherry-BMECs, cyan: siR-tubulin neuron label; scale bar, 500 µm), (iii) 3D integrated astrocyte and neuronal networks throughout the miBrain (green: Imaris surfaces of mCherry-astrocytes, cyan: siR-tubulin neuron label; scale bar, 500 µm), (iv) visualized at higher magnification (green: mCherry-astrocytes, cyan: tubulin neuron label, scale bar, 100 µm), and (v) anterior view of a miBrain 3D rendering (red: PECAM, cyan: neurofilament, blue: Hoechst; scale bar, 50 µm), (**K**) microglia: (i) 3D iMG distributed throughout BMEC networks throughout the miBrain (red: Imaris surfaces from mCherry-BMECs, green: Imaris surfaces from membrane pre-labeled iMG; scale bar, 500 µm), (ii) visualized at higher magnification (red: Imaris surfaces from mCherry-BMECs, green: Imaris surfaces from membrane pre-labeled iMG; scale bar, 100 µm), (iii) 3D iMG distributed throughout neuronal networks throughout the miBrain (cyan: tubulin neuron label, green: Imaris surfaces from membrane pre-labeled iMG; scale bar, 500 µm), (iv) visualized at higher magnification (cyan: Imaris reconstruction of tubulin neuron label, green: Imaris surfaces from membrane pre-labeled iMG; scale bar, 100 µm), and (v) distribution of iMG with BMEC and neuronal networks together (cyan: tubulin neuron label, red: Imaris surfaces from mCherry-BMECs, green: Imaris surfaces from membrane pre-labeled iMG; scale bar, 100 µm), (**L**) blood-brain barrier: (i) 3D astrocytes distributed throughout BMEC networks throughout the miBrain (red: Imaris surfaces from ZO1-BMECs, green: mCherry-astrocytes; scale bar, 500 µm), (ii) visualized at higher magnification (red: Imaris surfaces from ZO1-BMECs, green: Imaris surfaces from mCherry-astrocytes; scale bar, 100 µm), (iii) further magnified (scale bar, 50 µm), (iv) ZO-1 tight junctions along vessels (green: ZO-1, red: PECAM, blue: Hoechst; scale bar, 30 µm), (v) astrocytes with end-feet extending to vessels expressing canonical aquaporin-4 transporter (green: GFAP, gray: AQP4, red: PECAM, blue: Hoechst; scale bar, 30 µm; insert, gray: AQP4), and (vi) pericytes localized to the vessels (green: NG2, red: PECAM, blue: Hoechst; scale bar, 30 µm), and (**M**) myelinated neuronal networks: (i) 3D oligodendroglia distributed throughout neuronal networks throughout the miBrain (cyan: tubulin neuron label, green: Imaris surfaces from tdTomato pre-transfected oligodendroglia; scale bar, 500 µm), (ii) visualized at higher magnification (cyan: tubulin neuron label, green: tdTomato pre-transfected oligodendroglia; scale bar, 100 µm), (iii) myelin dye-labeled neurons and oligodendroglia (red: FluoroMyelin, cyan: tubulin neuron label, green: tdTomato pre-transfected oligodendroglia, scale bar, 100 µm), (iv) visualized also via Imaris reconstructions for myelin along with neurons and oligodendroglia (red: Imaris surfaces of FluoroMyelin, cyan: tubulin neuron label, green: Imaris surfaces of tdTomato pre-transfected oligodendroglia; scale bar, 100 µm), and (v) myelin and oligodendroglia alone (red: Imaris surfaces of FluoroMyelin, green: Imaris surfaces of tdTomato pre-transfected oligodendroglia; scale bar, 100 µm), and (vi) myelination of neuronal projections (green: MBP, cyan: neurofilament, blue: Hoechst; scale bar, 10 µm).

We utilized human brain data on cellular content to guide our construction approach. While the ratio between brain cell types has been debated over the last decades, more advanced methodologies suggest the glia-to-neuron ratio is less than 1-to-1^42^, with glia in the cortex estimated to be 45-75% for oligodendroglia, 19-40% for astrocytes, and 10% or less for microglia^43^. Single-cell RNA-sequencing (scRNAseq) approaches are providing further insights, though poor capture of vascular cell types and differential capture rates between cell types, could mask exact ratios. Endothelial cells are commonly estimated to comprise one-third of non-neuronal cells^44^. In practice, we found that incorporating cells at ratios of 39% neurons, 10% astrocytes, 6% oligodendroglia, 5% iMG, 39% BMECs, and 2% pericytes into miBrains formed an integral 3D tissue (**Fig. 1B**) with all of these cell types co-assembling together and distributing throughout the construct (**Fig. 1C**) in stable ratios, which we undertook to count for each cell type at weeks 1 and 2 (**Fig. S5A,B**). All cells are co-encapsulated into the Neuromatrix Hydrogel on day 0 with the exception of iMG, which are added to the co-culture on day 7 and migrated into the engineered tissue. Together the cells spread and formed 3D networks by day 5 and matured into an integral tissue mimic for the desired assay completion (**Fig. 1B,C; Fig. S5A,B**). High cell viability was observed throughout the cultures, including in neurons (**Fig. S5C,D**). In contrast to the emergent differentiation approach of organoids, in which days *in vitro* commence at the beginning of the differentiation, miBrains combine already differentiated and validated cells and we refer to their days *in vitro* beginning with their co-encapsulation together. The robust physiological features developed in the platform during the course of one to two weeks are a consequence of combining pre-differentiated cells in our optimized manner.

### Engineering a Brain Matrix-Inspired 3D Hydrogel

Current *in vitro* models typically co-culture a maximum of three different cell types stably. To enable the co-culture of all six major CNS cell types into an integral 3D tissue, we again referenced the human brain for guidance. The brain extracellular matrix (ECM) is predominantly comprised of a web of glycosaminoglycans and proteoglycans, enmeshing neuronal and glial cell types in the interstitial tissue, within which reside vascular networks ensheathed in a thin layer of BM proteins^45^. These ECM components provide both structural support and important biochemical signaling factors^45,46^.

We hypothesized that brain-matrix proteins normally expressed in the brain at later stages of development could be harnessed to accelerate maturation. We reasoned that if we could mimic brain tissue by constructing a soft 3D hydrogel with brain-mimetic components, we could provide scaffolding to encapsulate cells at the same time as promoting enhanced brain cell phenotypes. To rapidly assay differential effects of brain-matrix components we encapsulated combinations of hyaluronic acid, heparin, reelin, tenascin-R (TenR), tenascin-C (TenC), brevican (BCN), versican (VCN), neurocan (NCN), and thrombospondin-1 (TSP1) into standard Matrigel with BMECs, pericytes, astrocytes, oligodendrocytes, and neurons (**Fig. S6A**). The number of neuronal projections and presence of myelin basic protein (MBP) was low overall across these Matrigel-encapsulated conditions, with a small number of neuronal projections in the VCN condition (**Fig. S6A,B**). In parallel, we assayed electrical activity in these cultures via a multi-electrode array (MEA). Electrical activity was low across these conditions and only occasional spikes of neurons firing were detected even after several weeks in culture, though VCN again promoted a slightly increased firing rate (**Fig. S6C,D**). Activity in Matrigel was very low, with only occasional spikes (**Fig. S6C,D**).

As Matrigel is a heterogeneous admixture of mouse tumor BM proteins with limited mechanical and biochemical tunability^47^ and is rapidly degrading, as is common with protein-based biomaterials, we sought a more tunable and biochemically-defined scaffolding with which to present the brain-matrix cues. Dextran is a GAG biochemically similar yet distinct from hyaluronan, providing a uniquely brain-biosimilar scaffold, while still presenting a bioorthogonal backbone that can be mechanically tuned independently from biochemical cues^48^. Dextran hydrogels can be fabricated at soft stiffnesses and with viscoelastic properties akin to that of brain tissue^49^. We screened dextran-based hydrogels in conjunction with polyethylene glycol (PEG) hydrogels, a biochemically inert polymer that has similar advantages with respect to tunability, hyaluronan-based glycosil, and hyaluronan- and heparin-based heprasil. Encapsulating miBrain cells in each of these hydrogels in conjunction with crosslinker and BM peptide mimic, RGD, led to the establishment of a 3D co-culture inclusive of neuronal projections (**Fig. S6E,F**). Vascular networks were also observed in the dextran-based hydrogel conditions, while only small regions positive for BMEC marker PECAM were observed in other conditions (**Fig. S6E,F**). Further, on the MEA, miBrains encapsulated in PEG-, heprasil-, and glycosil-based hydrogels displayed only low levels of electrical activity (**Fig. S6G**). In light of these results, we decided to further screen dextran-based hydrogels, screening across mechanical compositions that could support integrated miBrain neurovascular unit formation. We cultured miBrains in dextran-based hydrogels across mechanical stiffness conditions by altering the molar amounts of dextran macromer and non-degradable peptide crosslinker, wherein intermediate molar ratios yielded the most robust vessel networks and neuronal projections (**Fig. S6H,I**). For this optimal stiffness condition, we modulated degradability by introducing a PEG non-degradable crosslinker at various concentrations. Both the 100% degradable and 90% degradable hydrogel promoted neurovascular unit assembly, with integrated microvascular networks and neuronal projections (**Fig. S6J,K**) and chose to utilize the 100% degradable condition to simplify the technical scale up.

Now with a brain-mimicking hydrogel scaffold in place to support miBrain co-culture, we revisited the hypothesis that brain matrix-specific proteins incorporated into a hydrogel, structurally capable of supporting their conjugation and presentation to cells, could be used to promote neuronal maturation. Screening across hydrogels incorporating HA, Heparin, Reelin, TenR, TenC, NCN, BCN, VCN, or TSP1, we co-cultured neurons and astrocytes and monitored neuronal activity on an MEA. After 2 weeks of culture, robust neuronal projections were observed across the hydrogel conditions, particularly in hydrogels incorporated with VCN, HA, TenC, and TSP1 (**Fig. S7A,B**). Strikingly, many of these matrix component conditions led to robust neuronal spiking on MEA recordings by week 2 in culture, a short onset time for iPSC-derived neurons. Representative raster plots of conditions demonstrate spiking across electrodes and bursts (**Fig. S7C**). Of the conditions tested, VCN, Reelin, BCN, TenC, NCN, and TSP1 displayed the highest firing rates (**Fig. S7D**). Given these promising results, we selected a subset of these ECM proteins that also supported hydrogel mechanical integrity and integrated these various brain-matrix proteins into our engineered dextran-based hydrogel with all miBrain cell types. Robust cultures and neuronal projections formed throughout the co-culture in many conditions, particularly for VCN (**Fig. 1D**). In terms of immunoreactivity of neuronal marker neurofilament, VCN and BCN led to the strongest signal intensity (**Fig. 1E**). Assessing these conditions on the MEA system in parallel, VCN and heparin displayed the highest neuronal firing rate (**Fig. 1F**). Across the many assays and experimental replicates tested, we found VCN to be a consistent top-performer. Our engineered dextran-based hydrogel incorporating VCN protein led to a reproducibly robust hydrogel scaffolding for the miBrain. While neurovascular units in miBrains suffered from the breakdown of Matrigel over time, robust neurovascular phenotypes persisted in our dextran-based hydrogel through the initial testing time of 5 weeks (**Fig. 1G**) and with enhanced structural integrity (**Fig. 1H**) and neuronal activity (**Fig. 1I**) compared to Matrigel. We therefore chose VCN-integrated dextran-based engineered hydrogels with RGD peptides and cell-degradable crosslinkers, terming the formulation “Neuromatrix Hydrogel.”

### 3D Immuno-Glial-Neurovascular miBrain Platform

To establish this novel miBrain platform as a tool that could be widely deployed to decode CNS disease mechanisms and accelerate drug discovery and development, we sought to characterize the cell- and tissue-scale biomimetic phenotypes constructed from our combination of six major CNS cell types together in our brain-mimicking 3D matrix. These features gave rise to 3D neurovascular units (**Fig. 1J, Movie S1**), throughout which were iMG resident immune cells (**Fig. 1K, Movie S1**), a biomimetic BBB (**Fig. 1L, Movie S2**), and myelin protein-positive neuronal networks (**Fig. 1M, Movie S3**), all co-integrated together in the miBrain. BMECs formed tubular networks while neurons formed networks of projections throughout the miBrain (**Fig. 1Ji,ii**). Astrocytes throughout the miBrain closely associated with neurons (**Fig. 1Jiii,iv**). The close proximity and cell-cell interactions between these cell types were observed at higher magnification, including between BMEC vessels and neuronal projections (**Fig. 1Jv**). iMG were dispersed throughout the miBrain tissue, including in close proximity to BMEC vessels (**Fig. 1Ki,ii**) and neurons (**Fig. 1Kiii,iv**), together in an integrated way (**Fig. 1Kv**). At the biomimetic BBB, astrocytes were proximal to BMEC microvessels throughout the miBrain (**Fig. 1Li-iii**), with BMECs expressing tight junctions (**Fig. 1Liv**), astrocytes expressing canonical transporter aquaporin-4 (**Fig. 1Lv**), and pericytes bordering the vessels (**Fig. 1Lvi**). Oligodendroglia closely associated with neurons throughout the miBrain (**Fig. 1Mi,ii**) and were surrounded by FluoroMyelin-positive signal, lining the neuronal networks (**Fig. 1Miii,iv**), which were also positive for myelin basic protein (**Fig. 1Mvi**).

### Characterization of Neuronal Phenotypes in miBrain

Evaluating neuronal markers via immunohistochemistry, miBrain-neurons were found to be largely MAP2-positive (**Fig. 2A**) and express synaptic proteins, such as the presynaptic vGlut1 and postsynaptic PSD95, and with a greater density than that of monocultured neurons (**Fig. 2B,C**). To assess neuronal activity in the miBrain we differentiated neurons from an iPSC line stably expressing tdTomato and GCaMP3. GCaMP-neurons integrated into miBrains formed projections and displayed expected neuronal morphologies. Live monitoring calcium transients in GCaMP-neurons, spikes of neuronal activity were recorded over time (**Fig. 2D**). Neurons in miBrains exhibited robust spikes of activity and displayed enhanced calcium transients compared to monocultured neurons (**Fig. 2E,F**). In parallel, we evaluated neuronal activity on the MEA system (**Fig. 2G**). Though miBrains are a 3D tissue and this method only captures activity towards the bottom surface, we were able to detect robust electrical activity (**Fig. 2H**). miBrains displayed enhanced spontaneous neuronal activity in terms of spike rate and burst rate (**Fig. 2I**) and promoted stimulus responsiveness, decreasing the latency time to spiking upon electrical stimulation and increasing the subsequently evoked spikes (**Fig. 2J**).

**Fig. 2:**
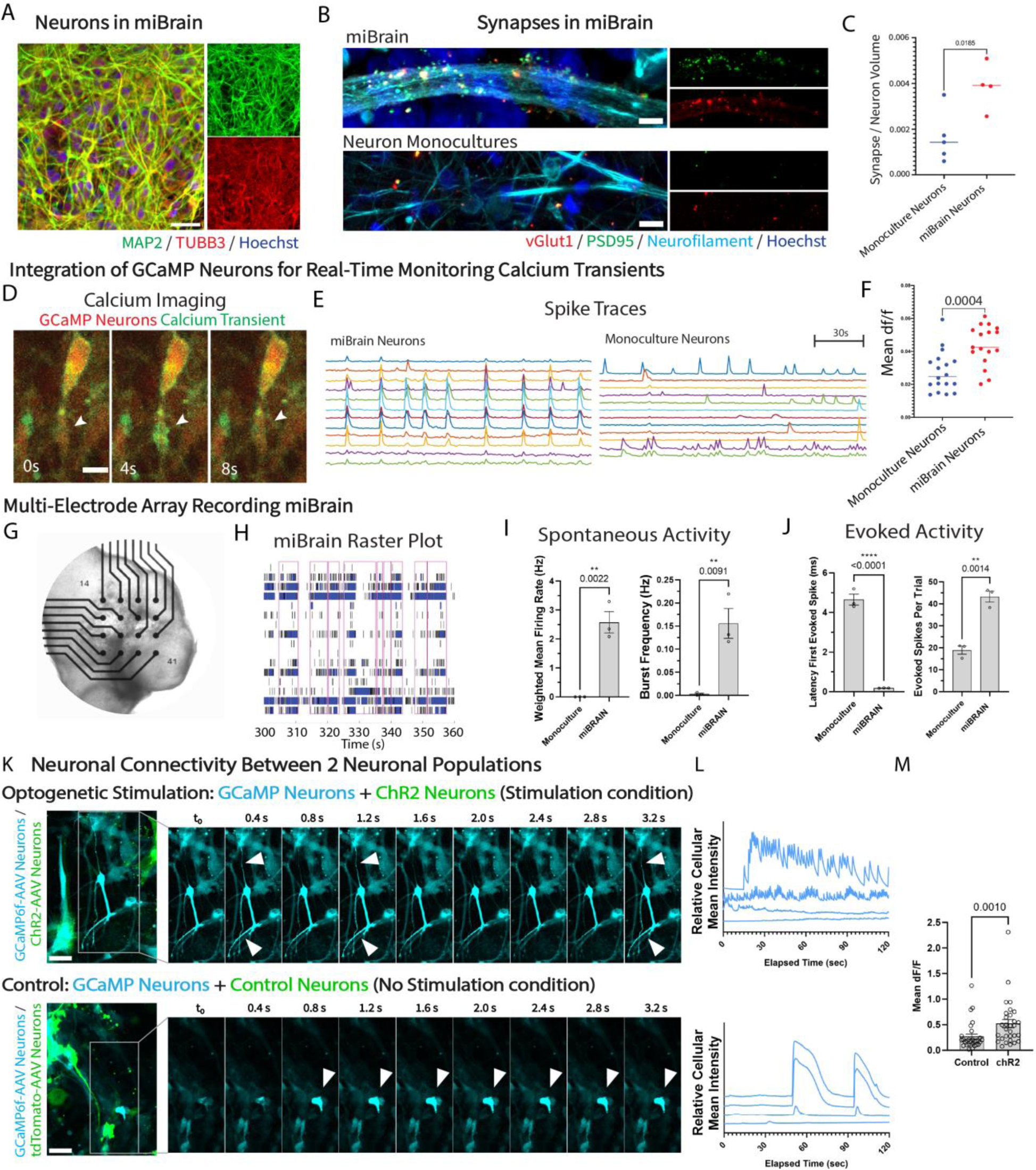
Neuronal phenotypes enabled by the miBrain. (**A**) Characterization of neurons integrated into miBrain immunohistochemistry (green: MAP2, red: TUBB3, blue: Hoechst; scale bar, 30 µm), (**B**) synapses in miBrain (top) versus neuronal monocultures (bottom) (red: vGlut1, green: PSD95, cyan: neurofilament, blue: Hoechst; scale bar, 5 µm), (**C**) quantification of synapses plotted as number of PSD95/synapsin co-localized puncta per volume neurofilament, n = 4, student’s t-test p = 0.0185), (**D**) example images of calcium dynamics in GCaMP neurons (red: tdTomato GCaMP neurons, green: calcium transient; scale bar, 25 µm), (**E**) example spike traces of neuronal calcium transients in miBrain (left) compared to monoculture (right) (scale bar, 30s), (**F**) quantification of calcium events plotted as max dF/F per calcium trace per neuron recorded (statistical analysis via student t test p = 0.0004), (**G**) macroscopic image of miBrain on MEA for monitoring electrical activity, (**H**) example raster plot miBrain at week 10, (**I**) characterization of spontaneous activity in miBrain compared to neuronal monocultures in terms of weighted mean firing rate (left) and burst frequency (right) (plotted as mean and S.E.M.; statistical analysis as unpaired t test), (**J**) evoked activity in miBrain compared to neuronal monocultures for latency after stimulation (left) and number of evoked spikes per trial (right) (plotted as mean and S.E.M.; statistical analysis as unpaired t test), (**K**) assessing neuronal connectivity between two neuronal populations in miBrain: neurons pre-transfected with GCaMP and a second neuronal population that is pre-transfected with either (top, left) ChR2-mCherry for optogenetic stimulation of one neuronal population under blue light, (cyan: GCaMP6f-neurons, green: ChR2-mCherry-neurons; scale bar, 30 µm), (top, right) displaying sequential images of calcium transients imaged under blue light (cyan: GCaMP6f-neurons), or (bottom, left) control tdTomato for the no stimulation condition (cyan: GCaMP6f-neurons, green: tdTomato-neurons; scale bar, 30 µm), displaying sequential images of calcium transients (cyan: GCaMP6f-neurons), (**L**) example spike traces of GCaMP6f-neurons in miBrains with a second population of neurons pre-transfected with (top) optogenetic ChR2-mCherry versus (bottom) control tdTomato, and (**M**) quantification of calcium events plotted as mean dF/F per calcium trace per neuron recorded (n = 3 miBrains/ group, repeated in 3 independent trials; statistical analysis via student t test p = 0.0010).

To evaluate the connectivity between neurons within the miBrain we incorporated two populations of neurons, between which activity was assessed with and without optogenetic stimulation. In both sets of miBrains, one population of neurons was pre-transfected with GCaMP6f via AAV. In one set of miBrains, a second neuron population was pre-transfected with Channelrhodpsin-2 (ChR2)-mCherry, while in the other miBrain set, the second neuron population was instead pre-transfected with a control AAV with tdTomato. This approach enables testing the responsiveness of one neuronal population to the heightened activity in another population, while visualizing these distinct neuronal populations. The excitation blue light for recording GCaMP6f-neurons, activates the ChR2-neurons in the stimulation condition and not the control neurons in the no stimulation condition (**Fig. 2K**). Increased spiking was observed in the GCaMP6f-neurons when co-cultured with the ChR2-neurons (**Fig. 2L**), with increased overall calcium transients, evidencing the connectivity between neuronal populations (**Fig. 2M**).

Assessing the responsiveness of miBrain-neurons to canonical channel modulators, we assembled a panel of pharmacological agents: voltage-gated sodium channel inhibitor TTX, AMPA competitive antagonist NBQX, NMDA receptor antagonist MK801, glutamate, which agonizes metabotropic glutamate receptors, and KCl, which triggers neuronal depolarization. Modulators were applied in increasing doses as neuronal activity was recorded on an MEA system. Doses of 8 nM TTX, 50 µM NBQX, and 50 µM MK801 decreased neuronal activity, assessed via mean firing rate, while 10 mM Glutamate and 1 mM KCl increased neuronal activity (**Fig. S8A**). Characterizing miBrain-neurons via whole-cell recordings, action potentials were recorded via current clamp (**Fig. S8B**) and inward and outward currents were recorded via voltage clamp (**Fig. S8C**). While establishing a patch-clamp within the 3D hydrogel required a non-ideal disruption of tissue integrity, we found miBrain-neurons displayed electrophysiological properties across these metrics, as well as for resting membrane potential (**Fig. S8D**) and input resistance (**Fig. S8E**) across samples (**Fig. S8F**).

To evaluate the transcriptomic changes that neurons undergo in the miBrain, we performed RNAseq (**Fig. S9A-C**). Monoculture- and miBrain-neurons differentially expressed a number of genes (**Fig. S9D**): neuronal genes were clearly upregulated in the miBrain condition (such as *SCN1A* and *CACNA1C;* **Fig. S9E**) and genes associated with neuronal precursor cells were downregulated (like *SOX11* and *PAX6;* **Fig. S9F**). These neuronal signatures provide evidence of strong neuronal phenotypes in the miBrain. Among the differentially expressed genes (DEGs) (**Fig. S9G,H**), genes associated with modulation of synaptic transmission, axon fasciculation, and astrocyte development are among those most strongly upregulated in miBrain-neurons. Conversely, neuronal stem cell population maintenance and neuroepithelial cell differentiation pathways are among those most strongly downregulated (**Fig. S9G,H**). In an effort to compare these datasets to human *in vivo* neurons, we correlated sequenced transcriptomes for miBrain- and monoculture-neurons to the whole genome of excitatory neurons in the prefrontal cortex from non-AD adult decedents^50^. When correlations to human *in vivo* excitatory neurons for miBrain-neurons and monoculture-neurons, differences were not observed when sampling over the whole genome (**Fig. S9Ii**), however, miBrain-neurons correlated more strongly than monoculture-neurons across pathways such as neuron differentiation (**Fig. S9Iii**) and axon ensheathment (**Fig. S9Iiii**).

### Characterization of Blood-Brain Barrier Integrated in miBrain

An ideal brain model would contain an integrated, 3D BBB expressing the physiologically-relevant transporters and receptors present in the human BBB, surrounded by the diversity of cell types that play key roles in reinforcing the barrier and contributing to disease pathogenesis. BMECs formed networks fully integrated into the miBrain and extending throughout the engineered tissue mimic (**Fig. 1C**). These microvascular networks formed 3D lumenized tubules expressing PECAM and tight junction markers ZO-1 (**Fig. 3A**) and CLDN-5 (**Fig. 3B**) and adhesion protein VE-CAD (**Fig. S10A**).

**Fig. 3:**
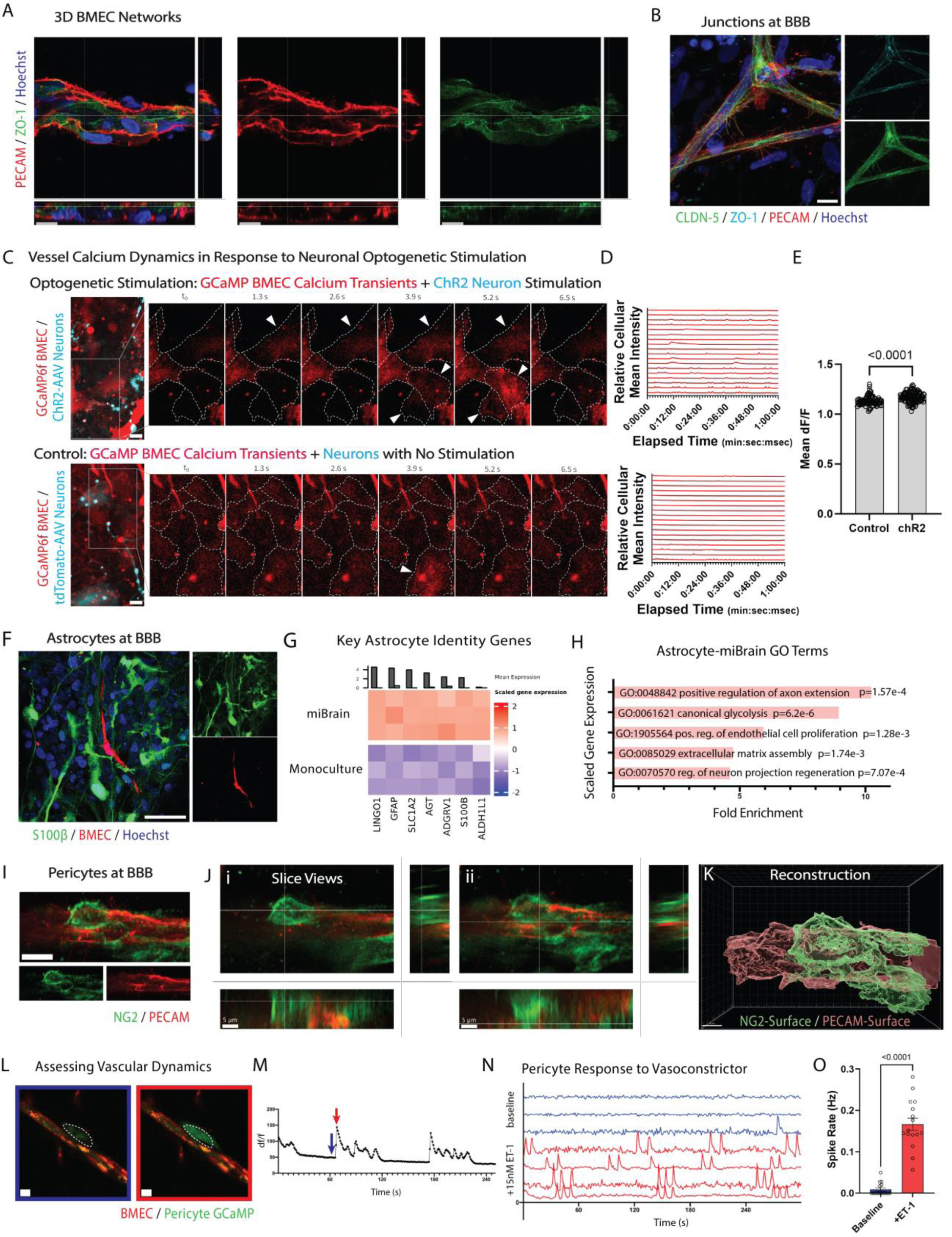
miBrain promotes brain-mimetic blood-brain barrier phenotypes. (**A**) Lumenized 3D BMEC vessels visualized in cross-sections of high magnification images (red: PECAM, green: ZO-1, blue: Hoechst; scale bar, 15 µm), (**B**) CLDN-5 tight junctions along vessels (green: CLDN-5, cyan: ZO-1, red: PECAM, blue: Hoechst; scale bar, 100 µm), (**C**) assessing neurovascular coupling via BMEC calcium transients in response to neuronal activity at baseline versus optogenetically-driven neuronal activity by incorporating GCaMP6f pre-transfected BMECs into miBrains with either (top, left) ChR2-mCherry-neurons for optogenetic stimulation of neurons under blue light, (red: GCaMP6f-BMECs, cyan: ChR2-mCherry-neurons; scale bar, 10 µm), (top, right) displaying sequential images of calcium transients imaged under blue light (red: GCaMP6f-BMECs), or (bottom, left) control tdTomato for the no stimulation condition (red: GCaMP6f-BMECs, cyan: tdTomato-neurons; scale bar, 10 µm), displaying sequential images of calcium transients (red: GCaMP6f-BMECs), (**D**) example spike traces of GCaMP6f-BMECs in miBrains with neurons pre-transfected with (top) optogenetic ChR2-mCherry versus (bottom) control tdTomato, and (**E**) quantification of calcium events plotted as mean dF/F per calcium trace per BMEC recorded (n = 3 miBrains/ group, repeated in 3 independent trials; statistical analysis via student t test p < 0.0001), (**F**) immunohistochemistry for astrocytes marker S100β (green: S100β, red: mCherry-BMEC, blue: Hoechst; scale bar, 50 µm), (**G**) expression of key *in vivo-*like genes in astrocytes isolated from miBrain compared to monocultured astrocytes from RNA-sequencing displayed as TMM-normalized and scaled expression for key astrocyte identity genes, (**H**) gene ontology analysis of astrocyte RNAseq for biological pathways upregulated in miBrain-cultured astrocytes based on significantly upregulated and downregulated DEGs and with an FDR p-value less than 0.05, (**I**) pericytes in miBrain localized to the microvascular networks (green: NG2, red: PECAM; scale bar, 10 µm), (**J**) visualized via slice and side views for z-planes (i) towards the top (superior) and (ii) middle (inferior) of the volume (green: NG2, red: PECAM; scale bars, 5 µm), and (**K**) corresponding 3D reconstruction (green: NG2-Surface, red: PECAM-Surface; scale bar, 4 µm), (**L**) calcium imaging of pericytes at vessels (left, blue) before and (right, red) during a signaling event (scale bars, 10 µm), (**M**) example traces of calcium transients, (**N**) spike traces (top, blue) with and (bottom, red) without application of vasoconstrictor ET-1, and (**O**) quantification of rate of calcium events (plotting mean and S.E.M., statistical analysis via paired t test).

To assess neurovascular integration of BMEC vessels and neurons we undertook an analogous approach to that used in assessing neuronal connectivity (**Fig. 2K**), in which we constructed miBrains with BMECs pre-transfected with GCaMP6f and either ChR2-mCherry-neurons or control tdTomato-neurons (**Fig. 3C**). BMECs in miBrains with control tdTomato-neurons displayed endogenous calcium transients. These BMEC calcium transients were more frequent in miBrains with optogenetically-stimulated ChR2-mCherry neurons (**Fig. 3D,E**). This indicates that BMECs in miBrains are responsive to enhanced neuronal firing and provides evidence of active neurovascular units in the miBrain.

Interacting with the brain microvasculature and forming the parenchymal niche of the BBB are astrocytes. These important glial cells, while also performing essential roles in neuronal regulation, have regulatory roles in vascular and barrier function^51^. iPSC-derived astrocytes, however, often have limited maturity with low expression levels of canonical fate markers. In the miBrain, astrocytes expressed canonical marker GFAP and water channel AQP4 and extended endfeet onto the vasculature (**Fig. S1Lv**). Further, miBrains harbored positive immunoreactivity for gap junction marker connexin-43 (Cx43) around the microvasculature (**Fig. S10B**) and canonical astrocyte marker S100β (**Fig. 3F**).

To characterize miBrain-astrocytes and better understand the effect of the miBrain niche on astrocyte signatures, we sequenced astrocytes (**Fig. S11A,B**). In the miBrain, astrocytes displayed a striking enrichment of identity genes like *S100β, SLC1A2,* and *ALDH1L1* (**Fig. 3G**), enhanced S100β (**Fig. S11H,I**) and GFAP (**Fig. S11J,K**) immunoreactivity, and a strong upregulation of a wide variety of genes involved in synaptic modulation and neuronal signaling (**Fig. S11C**) as well as vascular regulation (**Fig. S11D**). This points to the recapitulation of both barrier (including via *ROBO2* and *SEMA6A*) and neuronal process regulation (including via *NRP1*) in miBrain-astrocytes, among other DEGs (**Fig. S11E,F**). Consonantly, gene ontology analysis revealed a strong upregulation in biological pathways associated with neuronal processes and endothelial cell processes, as well as in extracellular matrix assembly (**Fig. 3H**). When these RNAseq datasets were compared to human *in vivo* astrocyte via a snRNAseq dataset from prefrontal cortex tissues of non-AD decedents^50^, miBrain-astrocytes correlated more strongly than monoculture-astrocytes (**Fig. S11G**).

While endothelial cells are the primary regulators of BBB permeability and selectivity, pericytes in the human brain are coupled to capillaries and exercise key functions in BBB maintenance, transport regulation, and cerebrovascular blood flow^52^. Correspondingly, in the miBrain pericytes localized to the outer lumen of the BMEC networks (**Fig. 3I-K**). As pericyte calcium increases in response to voltage-gated calcium channels and promotes contraction^52^, we sought to assess their functional coupling to the microvasculature. We constructed miBrains with pericytes stably expressing GCaMP6 and monitored their calcium transients in miBrains (**Fig. 3L,M**). We stimulated vessels with potent vasoconstrictor endothelin-1 (ET-1), known to induce a prolonged increase in blood pressure *in vivo,* and monitored pericyte calcium signaling as a proxy for pericyte contractile activity. ET-1 stimulation induced network-wide pericyte calcium transients and this elevated calcium activity persisted for minutes (**Fig. 3N,O**), indicating a robust pericyte response to vasomodulation.

BMECs, pericytes, and astrocytes together form the BBB, which is further modulated and influenced by the other cell types in the brain. To assess whether miBrain-BBBs respond to BBB modulators known to promote or reduce barrier function, we tested a panel of pharmacological agents. miBrains treated with thrombin exhibited decreased average vessel area and diminished total vessel area, while those treated with hydrocortisone displayed increased vessel area metrics (**Fig. S12A-C**). Dose curves of LPS and of TNF-α also led to smaller vessel areas and doses of 100 and 1000 ng/mL TNF-α also led to an overall decrease in total miBrain area (**Fig. S12A-C**). As ZO-1 is an important barrier marker, we assessed its expression across treatment conditions via immunohistochemistry. Consistent with expected results, we found that hydrocortisone enhanced vascular ZO-1 signal intensity while the other modulators decreased ZO-1 (**Fig. S12D,E**). To further assess changes in barrier function, we measured the membrane resistance of a BMEC monolayer with and without miBrain seeding proximal to the BMECs on a transwell system. BMECs with miBrains displayed increased resistance (**Fig. S10C**). Interestingly, comparing miBrains constructed with cells derived from an isogenic iPSC line CRISPR-edited to *APOE4/4* AD risk, BMEC monolayers seeded with these *APOE4/4* miBrains had decreased barrier function compared to *APOE3/3* miBrains as assessed via TEER on a transwell system (**Fig. S10C**).

### Characterization of Glial Phenotypes in miBrain

Microglia and oligodendrocytes regulate multiple processes in the brain, with key roles in neuronal health and function, while also contributing to vascular regulation and other physiological functions^53,54^. Through immunohistochemistry analysis, we observed that iMG incorporated into the miBrain integrated and extended processes in close association with the neurons and vasculature (**Fig. 4A**). miBrain-iMG displayed immunoreactivity to purinergic receptor P2RY12, involved in motility and associated with homeostatic processes (**Fig. 4A**).

**Fig. 4:**
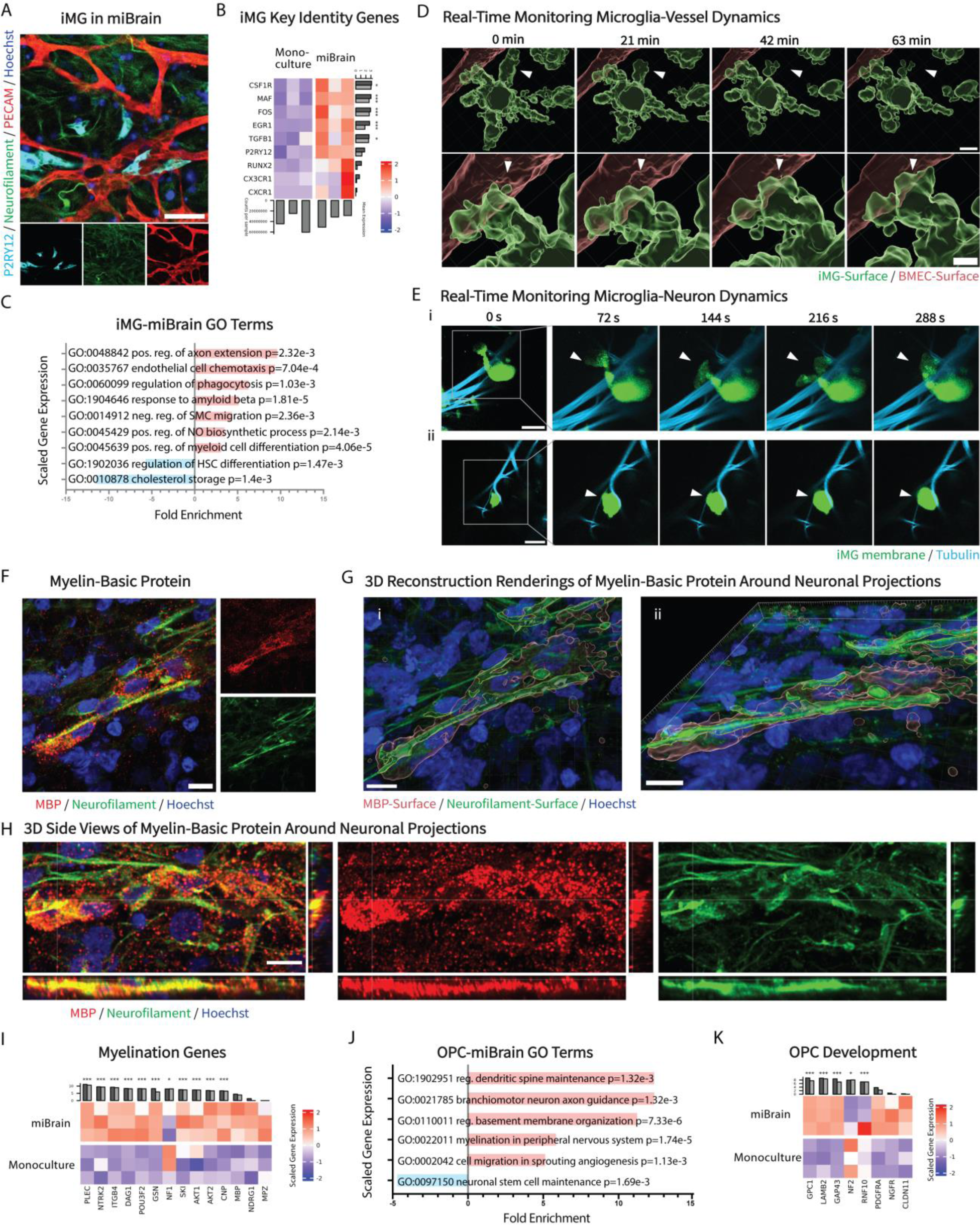
Glial phenotypes enabled by the miBrain. (**A**) Microglia-like cells in miBrain immunoreactivity to P2RY12 (cyan: P2RY12, green: neurofilament, red: PECAM, blue: Hoechst; scale bar, 50 µm), (**B**) expression of key *in vivo*-like genes in RNAseq of iMG isolated from miBrain compared to monocultured iMG displayed as TMM-normalized and scaled expression (* p < 0.05, ** p < 0.01, *** p < 0.001 for all RNAseq datasets), (**C**) gene ontology analysis of iMG RNAseq for biological pathways significantly altered in miBrain-cultured iMG based on significantly upregulated and downregulated DEGs and with an FDR p-value less than 0.05, (**D**) 3D, real-time monitoring of (top) iMG dynamics with vasculature in miBrain over 63 minutes (red: mCherry-BMECs-Surface, green: membrane pre-labeled iMG-Surface; scale bar, 10 µm) and (bottom) at higher magnification (scale bar, 5 µm), (**E**) real-time monitoring of iMG dynamics with neurons in miBrains over 5 minute recordings of (i) example 1 and (ii) example 2 (cyan: tubulin, green: GFP-membrane-labeled iMG; scale bar, 30 µm), (**F**) characterization of myelination produced by oligodendroglia in miBrain via immunohistochemistry (red: MBP, green: neurofilament, blue: Hoechst; scale bar, 10 µm), (**G**) 3D reconstructions of MBP-positive regions around neuronal projections viewed from (i) the top and (ii) the side (red: MBP-Surface, green: neurofilament-Surface, blue: Hoechst; scale bars, 10 µm), (**H**) side views of MBP-positive regions around neuronal projections (red: MBP, green: neurofilament, blue: Hoechst; scale bar, 10 µm), (**I**) expression of key genes associated with myelination in RNAseq of oligodendroglia isolated from miBrain compared to monocultured oligodendroglia displayed as TMM-normalized and scaled expression, (**J**) gene ontology analysis of oligodendroglia RNAseq for biological pathways significantly altered in miBrain-cultured oligodendroglia based on significantly upregulated and downregulated DEGs and with an FDR p-value less than 0.05, and (**K**) expression of key genes associated with OPC development in RNAseq of oligodendroglia isolated from miBrain compared to monocultured oligodendroglia displayed as TMM-normalized and scaled expression.

Assessing the transcriptional signatures of miBrain-iMGs, RNA isolated from iMG (**Fig. S13A,B**) revealed the miBrain leads to upregulated P2RY12 in iMG (**Fig. S13C**), which is also evident via flow cytometry assessment on the protein level (**Fig. S13D**). Between miBrain- and monoculture-iMG, a number of DEGs were identified (**Fig. S13E,F**). Importantly, key microglia identity genes were upregulated in the miBrain-iMG, including *CX3CR1, EGR1,* and *CSF1R* (**Fig. 3B, Fig. S5E**). Further, miBrain-iMG upregulated genes associated with neuronal signaling and endothelial cell processes (**Fig. 4C**). Comparing miBrain- and monoculture-iMG to a human *in vivo* dataset of adult non-AD prefrontal cortex tissue^50^, we found that miBrain-iMG correlated more strongly with human microglia (**Fig. S13G**). To better understand what pathways could contribute to this enrichment of biomimetic signatures, we further compared miBrain- and monoculture-iMG to human *in vivo* microglial states our laboratory has identified^55^. miBrain-iMG correlated more strongly to the homeostatic state than monoculture-iMG (**Fig. S13Hi**). Across the various states (**Fig. S13H**), the homeostatic state was the only one found to exhibit a statistically significant difference, though the neuronal surveillance state signature had the next highest p-value, increased in miBrain-iMG (p=0.06; **Fig. S13Hi**). These results indicate that the miBrain niche promotes enhanced microglia signatures and provides a system in which their multi-faceted interactions with vascular and neuronal components can be further probed in high-resolution and high throughput.

Given the important role of microglia in a variety of processes, including immune surveillance, blood-brain barrier function^56^, and neuronal activity modulation^57,58^, we sought to functionally monitor microglia dynamics in the miBrain in real-time. We pre-labeled iMG and incorporated them into miBrains with mCherry-labeled BMECs and tubulin-labeled neurons. We observed motile iMG migrating through the miBrain, including in close proximity to the microvasculature (**Fig. 4D**). Dynamic iMG were also observed closely associating with neurons (**Fig. 4Ei**) and physically interacting with neuronal projections (**Fig. 4Eii**).

Oligodendrocytes *in vivo* are the myelinating brain cell type that engage with neurons. Assessing oligodendroglia phenotypes in the miBrain, we detected positive myelin basic protein immunoreactivity (**Fig. 4F**) around neuronal projections (**Fig. 4G,H**). Analyzing the transcriptional signatures of oligodendroglia (**Fig. S14A-C**), miBrain-oligodendroglia displayed an upregulation of genes associated with myelination (**Fig. 4I,J**), biological pathways associated with dendritic spine maintenance and neuron axon guidance (**Fig. 4J**), including genes suggestive of neuronal interactions (such as *NDRG4, KIF1A,* and *SNAP25*; **Fig. S14D**) and of vascular interactions (such as *VEGFA, KDR, FLT4* and *NRP1*; **Fig. S14E**). Concomitantly, pathways associated with maintenance of neuronal stem cells (**Fig. 4J**) were strongly downregulated in miBrain-oligodendroglia while genes associated with maturation of OPCs were promoted (including *GPC1* and *PDGFRα*; **Fig. 4K**). miBrain-oligodendroglia displayed additional DEGs compared to monocultured oligodendroglia (**Fig. S14F,G**) and correlated more strongly to human *in vivo* oligodendroglia signatures identified via scRNAseq of prefrontal cortex, non-AD decedent brain tissue^50^ (**Fig. S14H**). Electron microscopy on miBrains further revealed ringed structures indicative of oligodendrocytes wrapping around neuronal axons (**Fig. S14I**).

### Harnessing miBrain to Investigate Functional Consequences of *APOE4* Astrocytes

The *E4* allele of the gene encoding lipid transport protein apolipoprotein E (APOE) strongly increases risk for developing late-onset AD and decreases age of disease onset compared to the *E3* allele^59^. Lipid accumulation in glia is a hallmark of AD^60^ and in *APOE4* astrocytes, a major lipid and cholesterol-producing cell type in the brain^61^, lipid metabolism pathways are perturbed^62^, lipids accumulate^63,64^, and cholesterol sequestration can promote brain matrisome changes^65^. The functional consequences of *APOE4* astrocyte dysfunction on the other cell types in the brain niche and the question of if they contribute directly to AD-associated pathologies remain poorly understood. We therefore sought to investigate phenotypic changes in the multicellular miBrain niche with *APOE4* astrocytes.

We constructed miBrains from isogenic sets of *APOE3/3* and *APOE4/4* iPSCs generated by CRIPSR-dCas9 editing^62^ and evaluated GFAP signal, given the increasing utilization of GFAP as a biomarker for AD^66,67^. We found that *APOE4* miBrains displayed increased GFAP intensity (**Fig. 5A,B**). Concomitantly, higher immunoreactivity to S100β, associated with neurodegenerative phenotypes^68,69^, was observed in the *APOE4* condition (**Fig. 5C,D**). While these differences were not appreciable in *APOE4* versus *APOE3* astrocytes in monoculture^62^ (**Fig. S15A**), these phenotypes were captured in the multicellular environment of the miBrain. miBrain-astrocytes are also responsive to insult, assessed via treatment with LPS, in which S100β (**Fig. S15B,C**) and GFAP (**Fig. S15D,E**) are further elevated compared to no-treatment controls. To evaluate if the results mirror expression of astrocytes in human brains, we analyzed a single-nucleus transcriptomic dataset for reactive astrocyte genes for *APOE4/4* and *APOE3/4* individuals versus *APOE3/3* individuals that our laboratory recently generated^20^. We found a broadly increased expression of reactive astrocyte genes in *APOE4* carriers compared to non-carriers, including for *GFAP, S100β,* and *VIM* (**Fig. 5E**). Among the panel of reactive astrocyte genes evaluated, the most strongly upregulated were *LINGO1, HSPB1,* and *CRYAB* (**Fig. 5E**). We therefore sought to assess the functional consequences of the increased reactive phenotype of *APOE4* astrocytes.

**Fig. 5:**
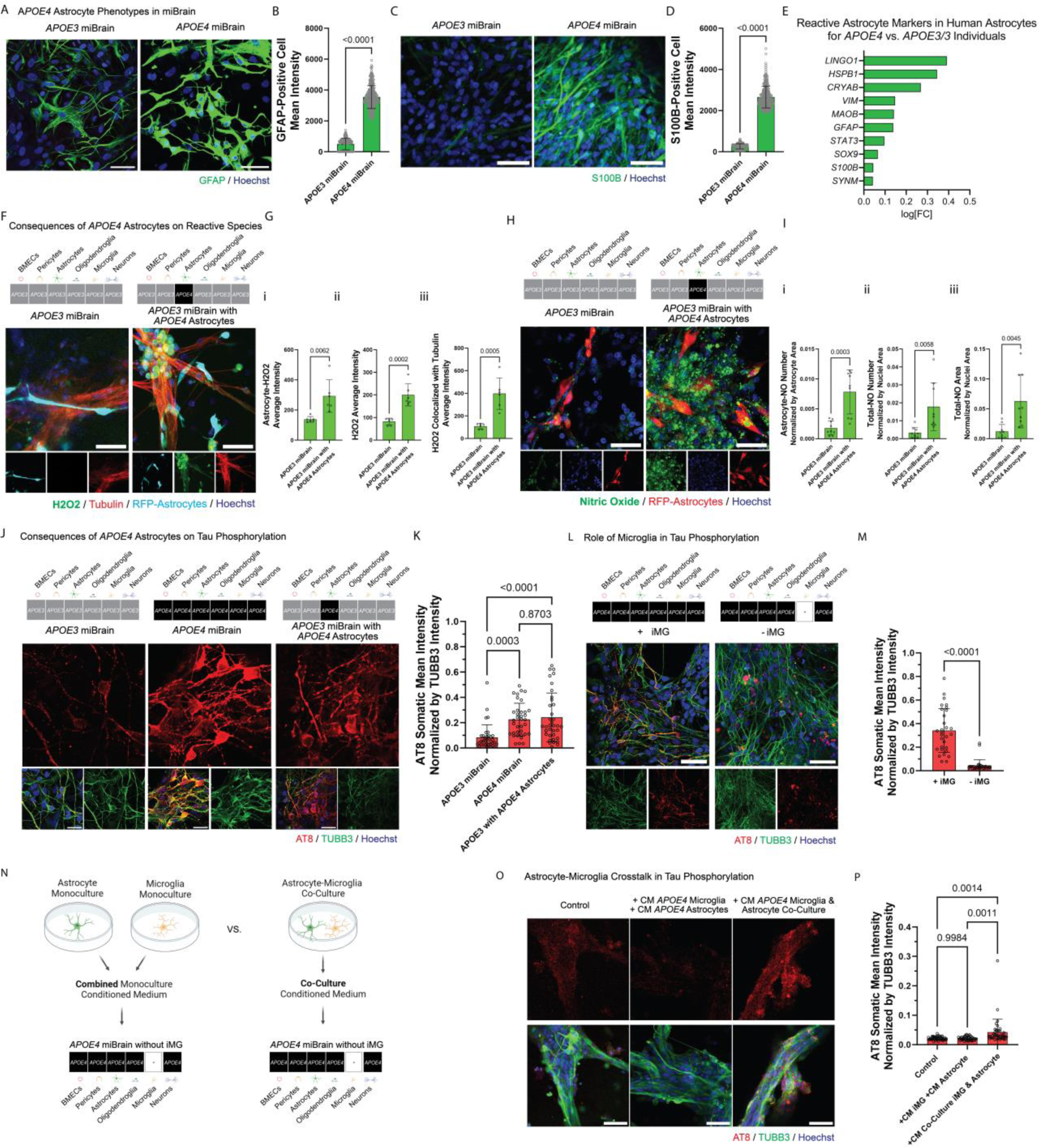
miBrain Reveals *APOE4* Astrocytes Promote Tau Phosphorylation via Crosstalk with Microglia. (**A**) GFAP immunoreactivity in *APOE3* (left) versus *APOE4* (right) miBrains (green: GFAP, blue: Hoechst; scale bars, 50 µm), (**B**) quantification of GFAP mean intensity normalized by nuclei area (n = 3 samples and images analyzed from n = 6 max projections from confocal z-stacks; statistical analysis by t test), (**C**) S100b immunoreactivity in *APOE3* (left) versus *APOE4* (right) miBrains (green: S100b, blue: Hoechst; scale bars, 50 µm), (**D**) quantification of S100b mean intensity normalized by nuclei area (n = 3 samples and images analyzed from n = 6 max projections from confocal z-stacks; statistical analysis by t test), (**E**) DEGs in human astrocytes from snRNAseq dataset^83^ of *APOE3/4* and *APOE4/4* versus *APOE3/3* individuals (positive log[FC] is upregulated in *APOE4*) for genes associated with reactive astrocytes, (**F**) hydrogen peroxide (H2O2) in miBrains with *APOE3* (left) versus *APOE4* (right) astrocytes incorporated into otherwise-*APOE3* miBrains (green: H2O2, red: live tubulin reagent, cyan: RFP-astrocytes, blue: Hoechst; scale bars, 50 µm), (**G**) quantification of hydrogen peroxide in terms of (**i**) mean hydrogen peroxide intensity co-localized with astrocyte region, (**ii**) mean hydrogen peroxide intensity in positive regions, and (**iii**) mean hydrogen peroxide intensity co-localized with tubulin-positive region (data from n = 3 replicates and images from n = 6 max projections from confocal z-stacks; statistical analysis via t test; experiment was conducted in 3 independent trials), (**H**) nitric oxide in miBrains with *APOE3* (left) versus *APOE4* (right) astrocytes incorporated into otherwise-*APOE3* miBrains (green: nitric oxide, red: RFP-astrocytes, blue: Hoechst; scale bars, 50 µm), (**I**) quantification of nitric oxide in terms of (**i**) nitric oxide segmented droplets co-localized with astrocytes normalized by astrocyte area, (**ii**) number of segmented nitric oxide (NO) droplets normalized by the nuclei area, and (**iii**) total area of nitric oxide segmented particles normalized by nuclei area (data from n = 3 replicates and images from n = 9 max projections from confocal z-stacks; statistical analysis via t test; experiment was conducted in 3 independent trials), (**J**) tau phosphorylation in *APOE3* miBrains (left), *APOE4* miBrains (middle), and *APOE3* miBrains with *APOE4* astrocytes (right) for miBrains treated with 20 nM exogenous Aβ 1-42 (red: AT8 phosphorylated tau, green: TUBB3, blue: Hoechst; scale bars, 30 µm), (**K**) quantification of tau phosphorylation assessed in the soma of TUBB3-positive neurons (mean intensity of AT8 normalized by mean intensity of TUBB3 per cell; data from n = 3 replicates and images from n = 12 fields of view; statistical analysis via ANOVA; experiment was conducted in 3 independent trials), (**L**) tau phosphorylation in *APOE4* miBrains with iMG (left) versus without iMG (right) for miBrains treated with 20 nM exogenous Aβ 1-42 (red: AT8 phosphorylated tau, green: TUBB3, blue: Hoechst; scale bars, 50 µm), (**M**) quantification of tau phosphorylation assessed in the soma of TUBB3-positive neurons (mean intensity of AT8 normalized by mean intensity of TUBB3 per cell; data from n = 3 replicates and images from n = 12 fields of view; statistical analysis via t test; experiment was conducted in 3 independent trials), (**N**) schematic illustrating experimental design for testing the effect of *APOE4* microglia-astrocyte crosstalk by collecting conditioned media from astrocyte monocultures and microglia monocultures (top) versus microglia-astrocyte co-culture (center) that is applied to *APOE4* miBrains in the absence of iMG (bottom), (**O**) tau phosphorylation in *APOE4* miBrain controls (left) versus miBrains treated with monoculture CM (middle) versus miBrains treated with microglia-astrocyte co-culture CM (right) co-administered with 20 nM exogenous Aβ 1-42 (red: AT8 phosphorylated tau, green: TUBB3, blue: Hoechst; scale bars, 50 µm), and (**P**) quantification of tau phosphorylation assessed in the soma of TUBB3-positive neurons (mean intensity of AT8 normalized by mean intensity of TUBB3 per cell; data from n = 3 replicates and images from n = 12 fields of view; statistical analysis via ANOVA).

While organoids cannot decouple astrocyte genotype from that of the other cell types due to their co-emergent differentiation from neuronal progenitors, the miBrain platform can decouple astrocyte genotype from that of the neurons and other cell types in the co-culture. Therefore, to assess astrocyte-specific contributions to *APOE4* pathogenesis we harnessed this advantageous miBrain capability to generate miBrains with *APOE3* neurons, iMG, oligodendroglia, pericytes, BMECs, and either *APOE3* or *APOE4* astrocytes.

Given that astrocytes play critical roles in regulating oxidative stress^70^ and that reactive astrocytes can aggravate neuronal damage^71^ and AD-associated pathological hallmarks, including via hydrogen peroxide production^72^, we examined reactive species in the miBrain. We detected higher levels of hydrogen peroxide colocalizing with astrocytes in the miBrains with *APOE4* astrocytes compared to the all-*APOE3* condition (**Fig. 5F,Gi**). Interestingly, increased hydrogen peroxide was also observed overall in the miBrains with *APOE4* astrocytes (**Fig. 5F,Gii**), including in neuron-colocalizing regions (**Fig. 5Giii**). Strikingly, bright intercellular droplets of nitric oxide were observed in miBrains with *APOE4* astrocytes (**Fig. 5H**). Between *APOE4-* versus *APOE3-* astrocyte miBrain conditions, more nitric oxide droplets were observed colocalizing with *APOE4* astrocytes (**Fig. 5H,Ii**) and in the overall for the *APOE4* astrocyte-containing miBrain, quantified in terms of total number and total area of nitric oxide droplets (**Fig. 5H,Iii,Iiii**). We similarly evaluated *APOE3* miBrains with *APOE4* versus *APOE3* astrocytes with a live indicator for peroxynitrite (**Fig. S15Fi**). While no differences were detected in astrocyte-colocalizing peroxynitrite (**Fig. S15Fii**), a significant overall increase in peroxynitrite species (**Fig. S15Fiii**) and neuron-colocalizing peroxynitrite (**Fig. S15Biv**), were detected. Together, these results provide evidence that not only do *APOE4* astrocytes harbor an increased proclivity to form hydrogen peroxide and nitric oxide species, but that they promote the accumulation of these species in neurons and other cells, which could contribute to *APOE4* neuropathologies.

Oxidative stress and free radicals, normally managed by lysosomal catabolism, can also, when imbalanced, contribute to lysosomal dysfunction, resulting in a vicious cycle that can exacerbate the cellular damage implicated in AD and other neurodegenerative diseases^73,74^. Evaluating lysosomal processes in our single-nucleus transcriptomic dataset of *APOE4* carriers and non-carriers^20^, we found differential gene expression in *APOE4* for genes associated with regulation of lysosomal catabolism, including *LRP1, LDLR,* and *ATP13A2* (**Fig. S15G**). Many related genes participate in regulation of oxidative stress-induced cell death, a pathway displaying upregulation of genes like *HSPB1, NFE2L2,* and *SOD1* and downregulation of *PRKN, PDE8A,* and *NME5* in astrocytes of *APOE4* carriers compared to non-carriers (**Fig. S15H**). Interestingly, miBrains with *APOE4* astrocytes compared to all-*APOE3* miBrains displayed a clear increase in lysosomal particle area colocalizing with astrocytes (**Fig. S15I,Ji**) as well as in overall lysosomal particle area (**Fig. S15Jii**) and lysosomal intensity (**Fig. S15Jiii**) in the miBrain. These data suggest that *APOE4* astrocytes both harbor lysosomal dysfunction and promote it in other cells, with this dysfunction correlating with the concomitant increase in reactive species observed in miBrains with *APOE4* astrocytes (**Fig. 5F-I**).

### miBrain Reveals *APOE4* Astrocyte Crosstalk with Microglia Promotes Neuronal Tau Phosphorylation

Given that the miBrain can uniquely dissect cell type-specific contributions to pathology with the construction of miBrains from cells of all decoupled genotypes, including that of astrocytes, we here sought to test whether *APOE4* astrocytes contribute to the *APOE4-*associated proclivity to tau phosphorylation. *APOE4* has been shown to accelerate amyloid aggregation^75^ and tau-mediated neurodegeneration^76^, and evidence suggests that removal of astrocytic APOE4 is protective against tau pathology^77^. Endogenously, *APOE4* miBrains displayed increased amyloid aggregation (**Fig. S16A,B**) and tau phosphorylation (**Fig. S16C-F**), here assessed at week 14 in culture, consistent with observations made by Lin et al., 2018^62^ using organoid culture. Notably, this includes the development of 4R tau pathology and at increased levels in *APOE4* compared to *APOE3* miBrains (**Fig. S16G,H**). To test whether *APOE4* astrocytes could promote tau phosphorylation, we compared *APOE3* miBrains with *APOE4* astrocytes to miBrains comprised of cells from all *APOE3* or *APOE4* backgrounds. To accelerate pathological progression, we administered a low 20 nM of exogenous amyloid beta peptide 1-42 (Aβ-42) resuspended at neutral pH. After 72 hours of treatment, increased tau phosphorylation was observed in *APOE4* miBrains via immunoreactivity to phosphorylated tau AT8, especially somatically (**Fig. 5J**). Interestingly, *APOE3* miBrains with *APOE4* astrocytes also displayed substantial tau phosphorylation and increased somatic AT8 intensity compared to *APOE3* miBrains (**Fig. 5J,K**).

Canonical reactive A1 astrocytes are known to be induced through the secretion of cytokines by microglia^78^ and microglia-astrocyte crosstalk is thought to mediate immune response to many neurological insult and disease conditions^79^. We thus asked whether microglia signaling modulates the *APOE4* astrocyte-associated tau phosphorylation we observed. We cultured miBrains with and without iMG. Strikingly, in the absence of microglia, tau phosphorylation was substantially diminished in *APOE4* miBrains (**Fig. 5L,M**). While some aggregates displayed strong immunoreactivity to phosphorylated tau, inter-neuronal tau colocalizing with tubulin marker TUBB3 was dramatically reduced (**Fig. 5L,M**). This suggests that microglia or microglia-derived cues are necessary to promote the observed tau phosphorylation in the context of *APOE4*.

To assess whether secreted factors resultant from microglia-astrocyte crosstalk contribute to the increased tau phosphorylation, we collected conditioned medium from *APOE4* microglia-astrocyte co-culture, treated with exogenous Aβ-42 as aforementioned, and applied it to *APOE4* miBrains (**Fig. 5N**). In parallel, we collected conditioned medium from monocultured *APOE4* astrocytes as well as from monocultured *APOE4* iMG, each similarly treated with Aβ-42, and pooled these media (**Fig. 5N**). Interestingly, the co-cultured microglia-astrocyte conditioned medium increased tau phosphorylation when applied to miBrains, while the combined monoculture conditioned media did not (**Fig. 5O,P**). This provides evidence that secreted factors from microglia-astrocyte signaling are sufficient to increase tau phosphorylation, even in the absence of microglia. The extent of tau phosphorylation observed in conditions of microglia-astrocyte co-culture applied to *APOE4* miBrains without iMG was less than that of *APOE4* miBrains with iMG (**Fig. 5L,M**), which could be due to the dose of soluble cues applied or suggest that other iMG-dependent processes contribute to the observed tau phosphorylation. However, the insufficiency of combined monoculture conditioned media compared to microglia-astrocyte co-culture conditioned media to promote tau phosphorylation, suggests that crosstalk between astrocytes and microglia produces factors distinct from their monocultured counterparts that promote neurotoxic phenotypes. These results could be in line with those of a recent study that reported that conditioned media from *APOE4* iMG with high lipid droplet burden, indicative of a pathological state, and not *APOE4* iMG with low levels of lipid droplets nor *APOE3* iMG, promoted tau phosphorylation in primary neuron culture^80^. In fact, examining lipid droplets in miBrains with *APOE3* versus *APOE4* astrocytes, we found increased lipid droplets in the *APOE4* condition, even in otherwise *APOE4* miBrain cell types (**Fig. S15K,L**). This evidences that *APOE4* astrocytes promote a pathological state in iMG that precipitates neurotoxic phenotypes.

## Discussion

We here describe a multicellular co-culture inclusive of all six major CNS cell types in an integral tissue mimic, to provide a model in which the contributions from each of these cell types can be queried. We engineered culture parameters to reconstruct important cell phenotypes, structural organization, and functional features of human brain tissue. We have established this miBrain platform mimicking the brain niche inclusive of a biomimetic BBB, neuronal networks, myelin-associated phenotypes, and immune cell components, with brain-mimetic architecture, hallmarks of brain physiology and pathology, and transcriptomic signatures. The Neuromatrix Hydrogel could be useful for other brain-modeling applications and the biomaterial design strategy underlying it could also inform the construction of vascularized tissues for other tissue types.

While this platform could provide substantial advantages in mechanistic understanding and disease modeling, it also has several limitations. We here use static culture and assess barrier function with junctional proteins as proxies, though future work can be aimed at introducing flow through the vessels within the miBrain via microfluidic approaches that could further enhance the platform biomimicry. Future work profiling neurons may be better performed using scRNAseq approaches that could more effectively isolate and capture neurons with the gentle droplet barcoding method. The viral transduction method used for neurons could also be less cell type-specific than the non-viral methods, which we were able to harness for isolating other miBrain cell types, in that other cell types like iMG could potentially phagocytose labeled cell debris, contaminating the virally-labeled population. Of note, while we here incorporated cells from the best iPSC differentiation protocols available, future iterations of the miBrain can integrate any advancements in the iPSC field, including in additional cell subtypes.

Harnessing this platform we have found that the miBrain niche can enhance iPSC-derived astrocyte signatures. We found that *APOE4* astrocytes could have an important role in mediating tau pathological progression, thus revealing pathological phenotypes conferred by non-cell autonomous interactions. This work has utilized the unique utility of our novel miBrain platform to recapitulate multicellular interactions in the CNS and assess cell type-specific phenotypes with decoupled genotypes. Advancing brain-mimicry *in vitro*, the miBrain platform could be a vehicle for accelerated mechanistic inquiry and therapeutic development for AD as well as other neurological indications. Scaling up the platform and miniaturizing the miBrain tissues, we have been able to robustly assess drug responses reproducibly in 48 and 96 well plate formats, positioning the technology for translation. This multicellular mimic inclusive of all six major brain cell types could provide advantages in dissecting disease mechanisms, modeling neuropathologies, and evaluating neurotherapeutics across indications.

## Materials and Methods

### Cell lines

All iPSC were maintained in mTeSR1 medium (Stem Cell Technologies) in feeder-free conditions. iPSC lines were previously generated by the Picower Institute for Learning and Memory iPSC Facility. iPSCs were passaged at 60-80% confluence using 0.5 mM EDTA for 5 minutes and reseeding 1:6 onto Matrigel-coated plates. Previously reported and characterized *APOE3/3-*parental iPSC line and its CRISPR-dCas9-edited isogenic *APOE4/4* line were used in this study^81^, along with corresponding NGN2-iPSCs^41^. The *APOE3/3* cell line was derived from a cognitively normal individual and edited to *APOE4/4* homozygous, while an *APOE4/4* line from an AD patient was corrected to *APOE3/3.* Results for *APOE4* astrocytes were corroborated with *APOE4/4*-parental and CRISPR-dCas9-edited isogenic *APOE3/3* line^41^. A ZO1-eGFP line (Allen Institute) was used for tracking experiments, as was a constitutive mCherry-expressing iPSC cell line, which was also used for RNAseq comparisons. For calcium imaging experiments comparing miBrain- and monocultured-neurons, tdTomato and GCaMP3 (addgene plasmid #62814) were introduced via the piggyBac transposon system into NGN2-iPSCs and successfully edited cells were purified based on tdTomato expression using flow cytometry to generate a stable line.

### Cell culture

BMECs were differentiated following a protocol similar to previously published protocols^29,30^, with modifications. In brief, on day 0, iPSCs were dissociated to a single-cell suspension and seeded onto basement membrane-grade Matrigel-coated plates at a density of 10,000 cells/cm^2^ in mTeSR1 supplemented with 10 µM ROCK inhibitor Y-27632. On day 1, medium was replaced with priming medium (DMEM/F12 media with GlutaMAX, 1% v/v MEM non-essential amino acids, and 1% v/v penicillin:streptomycin) supplemented with 6 µM CHIR-99021 and 10 ng/mL BMP4, feeding daily. On day 4, medium was replaced with endothelial induction medium (hECSR media, 2% v/v B27 supplement, 1% v/v MEM non-essential amino acids, and 1% v/v penicillin:streptomycin) supplemented with 50 ng/mL VEGF-A and 2 µM forskolin, feeding daily. On day 6, cells were passaged onto collagen-coated plates (50 µg/mL collagen type IV diluted in DPBS, coated for a minimum of 30 min at room temperature, then washed once with DPBS prior to use) at a split ratio of 1:3 in endothelial induction medium supplemented with 50 ng/mL VEGF-A and 2 µM forskolin, fed daily. On day 8, cells were passaged by incubating with StemPro Accutase for only ∼3-4 min to selectively dissociate endothelial cells, and replated onto collagen-coated plates at a split ratio of 1:5 in endothelial maintenance medium (hECSR media, 2% v/v B27 supplement, 1% v/v MEM non-essential amino acids, 1% v/v penicillin:streptomycin, 10 ng/mL VEGF-A). Cells were fed daily with endothelial maintenance medium and passaged as needed prior to reaching approximately 80% confluency to maintain endothelial cell character. After day 10, expression of endothelial cell-specific marker PECAM was assessed using flow cytometry prior to incorporation into the miBrain (**Fig. S2Ai**). Positive staining for canonical markers PECAM, vWF, CLDN-5, VE-CAD, and ZO-1 was confirmed via immunohistochemistry (**Fig. S1A**). Absence of pericyte marker PDGFRb and epithelial cell marker EPCAM, in addition to presence of microvascular marker CD34, were further confirmed by flow cytometry.

iPSCs were differentiated into pericytes as previously described^29^. In brief, iPSCs were maintained under feeder-free conditions until colonies reached approximately 60% confluence. On day 0, cells were dissociated using accutase and seeded on hES-grade Matrigel coated plates at a density of 3,000 cells/ cm^2^ and fed with mTeSR media supplemented with 10 μM ROCK inhibitor Y-27632. On day 1, media was replaced with priming media (DMEM/F12 media, 8 μM CHIR-99021, 25 ng/mL BMP4). On day 4, media was replaced with pericyte induction media (DMEM/F12 media, 100 ng/mL PDGF-B, 100 ng/mL activin A). On day 5, cells were passaged using accutase and replated on Matrigel coated plates at a density of 1e^5^ cells/ well of a 6-well plate and fed with pericyte induction media supplemented with 10 μM ROCK inhibitor. Pericytes were maintained without passage for an additional 14 days, feeding with pericyte expansion media (DMEM/F12 media, 10 ng/ml PDGF-B) every two days. Positive staining for canonical markers NG2 and PDGFRβ was confirmed via immunohistochemistry (**Fig. S2C**). Expression of pericyte-specific marker PDGFRβ was assessed using flow cytometry prior to incorporation into miBrains (**Fig. S2Aii**).

Astrocytes were generated according to a previously published protocol^33^, with modifications. Positive expression of astrocyte marker S100b and negative expression of neuronal progenitor marker SOX10 were validated using immunohistochemistry prior to incorporation into miBrain co-cultures (**Fig. S2D**). Expression of CD44 was assessed using flow cytometry (**Fig. S2Aiii**). OPCs were generated according to the protocol described in^31^. Positive expression of OPC markers PDGFRα and O4 were validated using FACS (**Fig. S2Aiv**) and immunohistochemistry (**Fig. S2E**) prior to incorporation into miBrain co-cultures. iMG were generated according a previously published protocol^34^. Positive expression of microglia marker CD45 was validated using FACS prior to incorporation into miBrain co-cultures (**Fig. S2Av**). Canonical marker detection via immunohistochemistry for Iba1, P2RY12, and TMEM119 was confirmed (**Fig. S2F**). Neurons were differentiated according to a previously published protocol^32^. In brief, iPSCs harboring tetON-NGN2 (NGN2-iPSC) were seeded on Matrigel-coated plates and treated with doxycycline (5μg/ml) for 5 days to induce NGN2 overexpression. Cells were treated with 5μM blasticidin for 3 days to eliminate populations lacking tetON-NGN2, then treated with 5μM puromycin for 2 days starting at day 3 to select for NGN2-positive cells. Neurons were dissociated and incorporated into miBrains on day 7 of this protocol. Expression of PSA NCAM was confirmed via flow cytometry (**Fig. S2Avi**) and positive marker expression of TUBB, MAP2, and NeuN was confirmed via immunohistochemistry (**Fig. S2G**).

Validated iPSC-differentiated cells were co-cultured in miBrains at the following ratios: 5 M/mL BMECs, 0.25 M/L pericytes, 1 M/mL astrocytes, 1 M/mL OPCs, 5 M/mL neurons, and 2.5 M/mL iMG. miBrains were cultured in Human Endothelial Serum-Free Medium (ThermoFisher Scientific) with 1% pen/strep (Sigma-Aldrich), 1% NEAA (ThermoFisher Scientific), 1% CD lipids (ThermoFisher Scientific), 1% Astrocyte Growth Supplement (ScienCell), 2% B-27 (ThermoFisher Scientific), 10 μL/ mL Insulin (ThermoFisher Scientific), 1 μM db-cAMP (Sigma Aldrich), 50 μg/ mL Ascorbic Acid (Sigma Aldrich), 10 ng/ mL NT-3 (PeproTech), 10 ng/ mL IGF-I (PeproTech), 100 ng/ mL Biotin (Sigma Aldrich), 60 ng/mL T3 (Sigma Aldrich), 4 μg/ mL Folic Acid (Sigma Aldrich) supplemented, 1 μM SAG (Tocris), and 50 ng/ mL VEGF-A (PeproTech). Upon the incorporation of iMG, 25 ng/mL M-CSF (Sigma Aldrich) was added and maintained in the media and after week 2, VEGF-A concentration was reduced to 10 ng/mL.

### Neuromatrix hydrogel encapsulation and miBrain formation

BMECs, pericytes, astrocytes, OPCs, and neurons were dissociated with accutase and counted using an automated cell counter (Countess). Cells were encapsulated in Neuromatrix hydrogel precursor solution adapting a dextran hydrogel protocol^82^, in which 3 mmol/L thiol-reactive groups of 2000 kDa Dextran-VS (Nanocs) or TRUE dextran (Sigma) were reacted with equimolar cell-degradable peptide (Sigma or GenScript), 3.0 mM CRGDS basement membrane-mimicking peptide (GenScript), and 50 μg/ mL human recombinant versican (R&D Systems), after bringing the solution to neutral pH. Cells resuspended in Neuromatrix were dispensed into the bottom of 48-well glass bottom plates (MatTek), allowed to polymerize at 37°C for 25 minutes, and then submerged in miBrain cell culture medium. The medium was exchanged after one hour to remove any reaction byproducts. miBrains were maintained in culture by feeding every other day. Experiments were performed between weeks 2 and 4 unless otherwise stated.

### Immunohistochemistry and confocal imaging

For immunohistochemistry on cell monocultures, cells were passaged onto Matrigel-coated chamber slides (MatTek) and fixed for 20 minutes in 4% PFA, washed with PBS, permeabilized in PBST (PBS, 0.1% Tween-20) for 10 minutes, incubated in blocking solution (PBS, 3% BSA, 2% donkey serum) for 30 minutes, incubated with primary antibody (1:500 unless otherwise specified) for 1 hour at room temperature or overnight at 4°C, washed five times with PBS and incubated with secondary antibody (1:500) and Hoechst 33342 (1:1000) in blocking solution for one hour at room temperature, then washed another five times.

For Claudin-5 BMEC immunohistochemistry, this protocol was followed, with the exception that cells were instead fixed in methanol by incubating for 5 minutes at -20°C, washing three times for 10 minutes with PBS, and then directly blocked in 2% serum and 3% BSA in PBS for 1 hour.

For miBrain immunohistochemistry, miBrains were fixed for 20 minutes in 4% PFA, washed with PBS, permeabilized in PBST (PBS, 0.1% Tween-20) for 30 minutes, incubated in blocking solution (PBS, 3% BSA, 2% donkey serum) for 2 hours, incubated with primary antibody (1:500 unless otherwise specified) overnight at 4°C, washed five times with PBS for a minimum of 15 minutes total and incubated with secondary antibody (1:500) and Hoechst 33342 (1:1000) in blocking solution for one hour at room temperature, and then washed another five times for a minimum of 15 minutes total.

High magnification imaging was performed using a Zeiss LSM 710 and LSM 900 confocal microscopes.

### Whole-miBrain imaging and cell number quantification

To image entire miBrains to visualize cell distribution and track miBrains over time, 9-by-9 tile scans were acquired over 264 μm z-height with a 20X objective in a Cytation 10 confocal imager system (Agilent BioTek). Images were stitched via the imaging processing tools in the imager software and z-projections generated via maximum intensity (BioTek).

To quantify cell numbers in the miBrain, 3-by-3 tile scans were acquired over 264 μm z-height with a 20X objective in a Cytation 10 confocal imager system at the center of the miBrain, stitched for each slice, and z-projections generated for maximum intensities (Agilent BioTek). Z-projections were input into Image J, a 3 sigma gaussian filter was applied to all images to minimize background, a consistent threshold was applied across images and timepoints for each cell type, the watershed function was applied to binary images, and the analyze particles function was applied for particles across a consistent size range for each cell type (pixel minimums set for each cell type. ZO-1 BMECs were used as the fluorescent ZO-1 junctions facilitated cell segmentation. Overlapping Hoechst stains were used to identify cells in the establishment of the protocol.

### Monoculture quantification

Non-junctional markers were quantified in Imaris following background subtraction of signal in iPSC controls stained in parallel for each marker of interest. Surfaces for Hoechst and the marker of interest were created for each image and the total number of overlapping cells was quantified.

BMEC junctional markers were quantified in ImageJ using the Trainable Weka Segmentation function to identify marker expression localized to cell borders. Briefly, WEKA classifiers were trained and applied on maximum intensity z-projections of confocal images for each marker across the entire image set. Reference channels (phalloidin or PECAM) were used to define cell borders via the Analyze Particles function (ROI size 5 µm – infinity to minimize noise). Identified border regions were overlaid on the junctional marker image and area of overlap was measured across all identified regions. Thresholds were set at 20% (CLDN-5 and VE-cad) and 2% (ZO-1) to determine if cells were positive for the junctional marker of interest based on manual confirmation on a smaller image set. This workflow is automated and unbiased in selecting border expression. However, the protocol is limited by the border-to-cytoplasmic signal ratio caused by background that can lead to an underrepresentation of the border signal and <100% cell confluency in samples that likely decreases junctional protein expression.

### TEER Assay

To assess trans-endothelial electrical resistance (TEER) across BMEC monolayers, 12 mm diameter Transwells with 0.4 µm Pore Polyester Membrane Inserts (CellQART) were coated with 400 µg/mL collagen I and 100 µg/mL fibronectin and seeded at 0.5 M cells/cm². When the cells reached 100% confluence, STX2 TEER Chopstick Electrodes (World Precision Instruments) were inserted with one electrode in the inner chamber of the Transwell and one electrode in the outer chamber of the Transwell and the measurements from a Millicell ERS2 Voltohmmeter (Millipore) were recorded. TEER was calculated by subtracting the resistance across a blank Transwell (no cells seeded on membrane, immersed in cell culture medium) and then multiplying by the membrane surface area.

### Live/Dead assay

To assess cell viability in miBrains, Calcein AM and Ethidium homodimer-1 from the LIVE/DEAD® Viability/Cytotoxicity Kit for mammalian cells were applied following manufacturer instructions (Invitrogen). For a reference control, one set of miBrains was pre-treated in 70% methanol for 30 min. In brief, cells were washed with DPBS, incubated for 30 minutes at room temperature while protecting from light with live/dead reagents and Hoechst nuclear label, and then imaged on a Cytation 10 confocal imager system with 4X objective and epifluorescence imaging mode, maintaining the same experimental protocol and imaging settings for each group (Agilent BioTek). To co-visualize neurons, miBrains were pre-incubated with a tubulin live dye overnight (Cytoskeleton) and also imaged in the far-red channel. Cell viability was quantified based on image intensity with the live fraction determined by the mean intensity of the live signal divided by the sum of live and dead mean intensities, applied across methanol controls and timepoint groups.

### 3D reconstructions and side views

Imaris software was used to visualize 3D z-stack images. Slice views were used to visualize lumenized vessels, pericyte localization proximal to vessels, and myelination around neuronal projections. Surfaces were computationally constructed from each imaging channel to visualize the distribution of cells throughout whole miBrains, pericyte localization proximal to vessels, iMG interactions with vessels, and myelination around neuronal projections.

### Antibodies

Primary antibodies for immunohistochemistry used are as follows unless otherwise specified:

| Target | Host | Vendor Information | Dilution |
| --- | --- | --- | --- |
| Neurofilament (smi311) | mouse | BioLegend 837802 | 1:500 |
| IBA1 | guinea pig | Synaptic Systems 234 004 | 1:500 |
| Tubulin (TUBB3, TUJ1) | rabbit, mouse | BioLegend 802001, 801213 | 1:500 |
| NG2 | rabbit | Abcam ab255811 | 1:500 |
| Aquaporin4 (AQP4) | rabbit | Thermo Fisher PA5-53234 | 1:500 |
| Myelin Basic Protein (MBP) | chicken | Millipore AB9348 | 1:500 |
| PECAM | sheep | R&D Systems AF806 | 1:500 |
| VE-Cadherin (VE-CAD) | goat | R&D Systems AF938 | 1:100 |
| ZO-1 | mouse | Thermo Fisher MA3-39100-A488 | 1:100 |
| Claudin 5 (CLDN-5) | mouse | Invitrogen 4C3C2 352588 | 1:250 |
| Von Willebrand Factor | rabbit | Abcam ab154193 | 1:100 |
| GFAP | goat | Abcam ab53554 | 1:500 |
| Connexin 43 (Cx43) | rabbit | Abcam ab11370 | 1:500 |
| S100β | rabbit | Abcam ab41548 | 1:500 |
| P2RY12 | rat | BioLegend 848002 | 1:500 |
| MAP2 | chicken | BioLegend 822501 | 1:500 |
| vGlut1 | chicken | Synaptic Systems 135316 | 1:500 |
| PSD95 | rabbit | Abcam ab18258 | 1:500 |
| Phospho-Tau (AT8) | mouse | Thermo Fisher MN1020 | 1:500 |
| PDGFRβ | mouse | Abcam ab69506 | 1:500 |
| SOX2 | goat | R&D Systems AF2018 | 1:500 |
| O4 | mouse | Millipore MAB345 | 1:500 |
| Synapsin | guinea pig | Synaptic Systems 106104 | 1:500 |
| Oct4 | rabbit | Cell Signaling Technology 2890 | 1:500 |
| Amyloid 82E1 | mouse | Immuno-Biological Laboratories 10323 | 1:500 |

Corresponding Alexa secondary antibodies (ThermoFisher) were used at 1:500. Alexa Fluor 647 phalloidin (1:400) was incubated along with secondary antibodies.

### Flow cytometry

Cells were dissociated using accutase and washed with FACS buffer (DPBS, 0.5% BSA, 2 mM EDTA), filtered through a 40 μm cell strainer, then resuspended in blocking buffer (DPBS, 1% FBS) with 1:200 antibody and incubated for 25 minutes on ice protected from light. Cells were then washed with FACS buffer and resuspended in buffer with DAPI before filtering through a 40 μm cell strainer and proceeding with FACS. Flow cytometry was performed on a BD LSRFortessa cell analyzer equipped with 355 nm, 405 nm, 488 nm, 561 nm, and 640 nm lasers. Positive expression of cell type specific markers was assessed relative to isotype controls.

Antibodies used for flow cytometry were as follows:

| Target | Host / Isotype | Vendor Information | Dilution |
| --- | --- | --- | --- |
| CD31-FITC | Mouse IgG1 | BioLegend 303104 | 1:200 |
| CD140b-PE | Mouse IgG2a | BD 558821 | 1:200 |
| CD44-APC | Mouse IgG1 | BioLegend 338806 | 1:200 |
| PDGFRa-APC | Mouse IgG1 | R&D systems FAB1264A | 1:200 |
| PSA NCAM-APC | Mouse IgM | Miltenyi Biotec 130-120-437 | 1:200 |
| CD45-APC/Cy7 | Mouse IgG1 | BD 561863 | 1:200 |
| FITC mouse IgG1, k isotype control | Mouse IgG1 | BioLegend 400107 | 1:200 |
| PE mouse IgG2a, k isotype control | Mouse IgG2a | BD 555574 | 1:200 |
| APC mouse IgG1, k isotype control | Mouse IgG1 | BioLegend 400119 | 1:200 |
| APC mouse IgM isotype control | Mouse IgM | Miltenyi Biotec 130-122-931 | 1:200 |
| APC/Cy7 mouse IgG1, k isotype control | Mouse IgG1 | BD 557873 | 1:200 |
| P2RY12-PE | Mouse IgG2a | BioLegend 392103 | 1:200 |
| PE mouse IgG2a, k isotype control | Mouse IgG2a | BioLegend 400213 | 1:200 |

### Isolation and RNAseq of cell types from miBrain

To isolate cells from miBrains for RNAseq, miBrains were matured for 2 weeks prior to cell isolation and RNA sequencing. Corresponding cell monocultures were maintained in parallel for 2 weeks and then sorted using the same parameters as the miBrains. To minimize batch-dependent effects, the same batch of differentiated cells was used for the monoculture condition and the miBrain condition in each RNAseq experiment.

Neurons were isolated by first virally labeling neurons from the same batch with pAAV-CAG-tdTomato for 24 hours (1:50 dilution, Addgene 59462-AAV1) prior to incorporation into miBrain or monoculture, a subset of which were incorporated into miBrains and a subset of which were maintained in monoculture. To ensure sufficient capture of target cell populations, 15 individual miBrains were pooled per replicate, and three replicates were isolated per experiment. After 2 weeks, tdTomato-neuron miBrains and monoculture samples were harvested. Astrocytes and oligodendroglia were isolated using a similar approach, but by utilizing endogenously mCherry-expressing astrocytes. iMG were incorporated into 2-week old miBrains, while a subset from the same batch were maintained in monoculture, and after an additional four days, sorted based on CD45-Cy7.

To dissociate miBrains, samples were washed with PBS, then were incubated for 5-10 minutes in a mixture of accutase, collagenase IV (1:20), and dextranase (1:20). These were triturated 15-20 times with a P1000 pipette to form a homogeneous suspension, then pelleted and washed with DPBS twice. Astrocytes, iMG, and neuron monocultures grown in parallel were dissociated using accutase, then washed with DPBS prior to proceeding with antibody labeling and sorting. Oligodendroglia were fixed with 4% PFA on ice for 20 minutes prior to dissociation to ensure stability. iMG were incubated with CD45-Cy7 alongside isotype controls at 1:200 on ice for 25 minutes before proceeding with sorting.

Dissociated miBrains and monocultures were resuspended in FACS buffer (DPBS, BSA, EDTA) with DAPI and filtered through a 40 μm strainer. Samples were sorted on a BD FACSAria equipped with a 70 μm nozzle. Neurons were FACS-isolated based on tdTomato expression, astrocytes and oligodendroglia based on mCherry, and iMG based on CD45 expression. RNA was harvested in triplicate and in parallel from 20K neurons, 50K astrocytes, 50K oligodendroglia, or 2K CD45-positive iMG per replicate. Cells were gated on FSC-A vs SSC-A to separate intact cells from debris, then FSC-H vs FSC-W and SSC-H vs SSC-W to separate single cells from doublets, then DAPI to exclude dead cells, and then the channel corresponding to the antibody used. Cells were sorted into 96-well plates containing 50 μl RNAprotect per well.

For live samples, RNA was isolated from sorted cells using the Qiagen RNeasy isolation kit according to the manufacturer’s protocol. For fixed samples, sorted cells were spun down and resuspended in 200 μL digestion buffer (RecoverAll kit) supplemented with 1:50 Proteinase K, incubated at 50°C for 15 minutes, and then at 80°C for 15 minutes with no agitation to de-crosslink RNA from fixed tissue. 800 μL Trizol was added to the sample and incubated for 5 minutes, then 215 μl chloroform was added and the sample was vortexed vigorously for 30 seconds. Sample was transferred to a 5-prime phase lock heavy gel tube and centrifuged for 15 minutes at 12,000 g. The upper layer containing RNA was transferred to a fresh 1.5 ml microcentrifuge tube and an equal volume of 100% ethanol before proceeding with RNA isolation using Zymo Direct-zol RNA microprep kit according to the manufacturer’s protocols. Standard RNA quality control was performed using AATI fragment analyzer. RNA sequencing libraries were prepared using NEB Ultra II PolyA stranded library and sequenced using Illumina NextSeq500.

### RNA-sequencing data analysis

RNA-sequencing analysis was performed for each batch in a similar fashion. For preliminary QC, FastQC was used to look at quality. For each sample, gene reads were mapped to GRCh38.p13 and genes to GENCODE v43 using STAR 2.7.9a. These were then counted as reads per-sample using featureCounts from Subread version 1.6.2. Each sample was loaded into R (version 4.2.2) using a BioConductor (version 3.16) based workflow to perform QC and further analysis. Reads were normalized to trimmed mean of M-values (TMM) normalization using the R command “cpm(calcNormFactors(rawCounts, “TMM”))” in edgeR. Differential expression analysis was also performed using edgeR, keeping genes with a threshold of 100 counts per million (CPM) in at least 25% of the samples used in differential expression analysis. The DEGs were calculated in edgeR using a quasi-likelihood F-test with no covariates. Comparisons using a likelihood ratio test (LRT) in edgeR and DESeq2 were performed; but similar results were obtained. Gene ontology analysis was performed on significantly upregulated and downregulated DEGs using Panther DB, which identified GO terms with an FDR p-value less than 0.05. From these GO terms were selected the most biologically salient pathways, presented in order of fold-enrichment. iPSC data was downloaded and reprocessed using the same above RNA-sequencing workflow from GEO GSE102956. Single-cell RNAseq data for comparisons was downloaded from the Mathys, et al., 2023^50^ aging brain atlas from the linked website https://compbio.mit.edu/ad_aging_brain/, filtered for non-pathological samples, and summarized to major cell type per-donor by summing gene counts per sample. Microglia states were similarly processed from https://compbio.mit.edu/microglia_states/^55^. Comparisons between bulk RNAseq were performed by taking common genes between the two references above 100 counts per million in 25% of each of the groups across each RNAseq sample group in order to make the comparisons the same across case and control. Code utilized to analyze the data is included on GitHub at https://github.com/kellislab/miBrain. PCA gene loadings of BMECs to prior literature were performed by running PCA on the Z-scored TMM expression of samples and taking the top 100 gene loadings of the first principal component and the bottom 100 gene loadings of the first principal component.

### Calcium imaging of neurons

To generate neurons with GCaMP expression, iPSCs containing tetON Ngn2 were transfected with a plasmid driving GCaMP3 expression under the CAG promoter (Addgene #62810), then differentiated into neurons and incorporated into miBrain cultures as described above. In order to facilitate live imaging, miBrains were cultured on 3-cm glass bottom dishes (MatTek). Two-photon microscopy was performed on an Olympus FVMPE-RS microscope equipped with SpectraPhysics InsightX3 DeepSee laser tuned to 920nm. Images were acquired using Fluoview acquisition software (Olympus), we simultaneously acquired images in the red channel (bandpass filter 575-645 nm) to visualize tdTomato-expressing neurons, and the green channel (bandpass filter 495-540 nm) for GCaMP signal. Time courses of spontaneous calcium activity were collected at various time points, starting at week 4. Regions of interest were defined as tdTomato-positive neuronal soma and were manually outlined using ImageJ software and then quantified in Matlab. Calcium spikes were defined as periods in which GCaMP signal exceeded two standard deviations above baseline.

### Optogenetic evaluation of neuronal connectivity and neurovascular responsiveness

NGN2-neurons were differentiated from NGN2-iPSCs through the 7-day protocol and then transfected with either pAAV-CAG-hChR2(H134R)-mCherry (Addgene) or pAAV-CAG-tdTomato (Addgene). To evaluate the connectivity of miBrain-neurons, a second population of neurons were transfected with pAAV-CAG-GCaMP6f (Addgene). To evaluate vascular response to neuronal activity, BMECs were transfected with pAAV-CAG-GCaMP6f (Addgene). 48 hours later, positive transfection was confirmed, transfected cells were rinsed with PBS three times prior to dissociation, and miBrains were constructed. To evaluate neuronal connectivity, one set of miBrains was constructed with equal numbers of GCaMP6f- and ChR2-neurons, while a second control set was constructed with equal numbers of GCaMP6f- and tdTomato-neurons. To evaluate vascular response to neuronal activity, one set of miBrains was constructed with ChR2-neurons and GCaMP6f-BMECs, while a second set was constructed with tdTomato-neurons and GCaMP6f-BMECs. Calcium transients were recorded at day 10 using a Cytation 10 confocal imaging system (Agilent BioTek) with a 0.08 s sampling rate for neuronal recordings and 0.13 s sampling rate for BMEC recordings, imaging GFP with blue excitation light.

### Calcium imaging of pericytes

To generate pericytes with GCaMP expression, iPSCs were transfected with a lentivirus driving GCaMP6 expression under the CAG promoter, then differentiated into pericytes and incorporated into miBrain cultures together with BMECs generated from an iPSC line with constitutive mCherry expression. Calcium imaging experiments were performed on 2-week old miBrains using the Olympus FVMPE-RS 2-photon microscope as described in the previous section. For vascular dynamics assay, spontaneous activity of pericytes was recorded for approximately 5 minutes, then 100 nM ET-1 was manually injected, followed by a recovery time of 5 minutes before imaging calcium activity for 5 minutes. Regions of interest were defined as pericyte cell bodies and manually outlined using ImageJ software. A calcium transient was defined as a period of time in which mean pericyte calcium signal intensity was greater than two standard deviations above the mean of baseline.

### Calcium imaging of astrocytes

To characterize calcium activity in iPSC-derived astrocytes, astrocytes at the conclusion of the differentiation protocol were re-plated in an assay plate at 100 K/cm^2^ in astrocyte medium and allowed to recover for 2 days. Astrocytes were rinsed with HBSS and 30 minutes later, Fluo-4, AM (ThermoFisher Scientific) was applied at 1:1000 in astrocyte medium and incubated for an additional 30 minutes. Recordings were collected using the Cytation 10 automated imager system (Agilent BioTek) with environmental control.

### MEA recording

A MaestroPro MEA system (Axion Biosystems) was utilized to record electrical activity on miBrains. miBrains were seeded directly onto MEA plates (Axion Biosystems) and maintained in culture according to our standard protocol. 30 minute recordings were collected from miBrains and analyzed using Neural Metrix software (Axion Biosystems).

To assess the effect of neuromodulators on miBrain-neuron activity, activity was first recorded on the MEA system with 5 min. recordings. Tetrodotoxin (TTX; Tocris; 8, 80, 800 nM), NBQX (Tocris; 1, 10, 50, 200 μM), (+)-MK 801 maleate (MK801; R&D Systems; 50, 100, 500 μM), MNI-caged-L-Glutamate (R&D Systems; 0.1, 1, 10, 20 mM), and KCl (Sigma-Aldrich; 0.5, 1 mM) were applied and activity was recorded 2 minutes after dosing (n = 4 miBrains/ group). The fold change in mean firing rate from before versus after dosing was calculated for each treatment condition.

### Assessment of BBB Modulators on Barrier Function

miBrains were constructed with endogenously mCherry-expressing BMECs. On day 4, thrombin from bovine plasma (Sigma-Aldrich; 100 nM), hydrocortisone (Sigma-Aldrich; 140 nM), lipopolysaccharides (LPS) from *Escherichia coli* O26:B6 (Sigma-Aldrich; 10, 100, and 1000 μg/mL), and recombinant TNF-α (Peptrotech;10, 100, and 1000 ng/mL) were applied. Whole-miBrain images were collected after 24 hours (Cytation 10, Agilent BioTek, 4x objective, z-stack tile-scans). In Fiji, vessels were analyzed via automated analysis, in which the analyze particles function was applied to maximum projection images of whole-miBrains to analyze vessels of 100 square μm and above. The average vessel area and total vessel area were recorded and plotted in GraphPad Prism. Whole-miBrain areas were quantified via automated analysis from the vascular regions, increasing the threshold to include all area within the vessel networks as positive area. miBrains were fixed at 48 hours, immunolabeled for PECAM and ZO-1, and imaged via confocal microscopy (Cytation 10, Agilent BioTek, 20x objective, z-stack tile-scans). In Fiji, vessel regions were defined using the PECAM channel via automated analysis, ZO-1 signal intensity was assessed within PECAM-positive regions, and results were plotted in GraphPad Prism.

### Assessment of Astrocyte Response to LPS treatment

miBrains were matured for 1 week before applying 100 µg/mL LPS for 48 hours versus no-treatment controls (n = 4 miBrains/ group). After 48 hours, miBrains were fixed in 4% PFA and analyzed via immunohistochemistry.

### Amyloid treatment

Stock solutions of amyloid beta 1-42 peptide (Anaspec) were resuspended in PBS. Peptides were administered to miBrains at 20 nM for three days to induce pathology.

### Real-time monitoring miBrain

Live imaging of miBrains were performed on a Zeiss LSM 900 confocal microscope using a 20x objective with environmental control of humidity, temperature, and CO2. Reactive species were assessed by treating miBrains with biosensors for nitric oxide (siRNO Nitric Oxide Live Cell Dye, Sigma-Aldrich), peroxynitrite (BioTrack 515 Green ONOO-, Sigma-Aldrich), or hydrogen peroxide (Green H2O2, Sigma-Aldrich) following the manufacturer instructions. Mitochondria were similarly assessed using BioTrack 633 Red Mito Dye (Sigma-Aldrich) and lysosomes using BioTrack NIR633 LysO Dye (Sigma-Aldrich). Tubulin was similarly visualized live with a small molecule dye (Cytoskeleton). Astrocytes were labeled for live assays prior to incorporation into miBrains using an AAV (pAAV-CAG-tdTomato, Addgene or AAV9-CAG-GFP, Applied Biological Materials). Microglia were labeled for live assays prior to incorporation using a cell membrane dye (Vybrant Cell-Labeling Solutions, Life Technologies). A stably expressing mCherry cell line was utilized for live imaging endothelial cells that was previously developed using pAAVS1-P-CAG-mCh (Addgene). FluoroMyelin dye was used for live labeling myelin in miBrains (Thermo Fisher Scientific).

### Western blots

Whole tissue lysate protein fractions from miBrains were prepared by washing samples with cold PBS, then adding RIPA buffer with protease inhibitor (150mL/sample) (Sigma-Aldrich cat: R0278) and incubating on ice for 20 minutes. Tissue was manually disrupted with a p1000 pipette, then centrifuged at 14,000g for 15 minutes at 4°C. Supernatant containing the protein fraction was then saved and quantified using a Pierce 660 assay (Thermo Fisher cat: 22662). For western analysis, 10 μg of total protein in 6X Laemmli SDS sample buffer (Thermo Fisher Scientific) was heated for 5 minutes at 95°C and subjected to SDS-PAGE. Electrophoresis and transfer onto PVDF membranes was performed using the Biorad Mini-Protean Gel Electrophoresis System and Trans-blot Turbo Transfer System, according to manufacturer recommendations. After transfer, membranes were blocked in TBST (Tris-buffered saline, 0.02% Tween-20) with 5% BSA for 1 hour, and incubated overnight with primary antibody at 4°C in a 1:1000 dilution. After subsequent washes with TBST, membrane was incubated in HRP-linked secondary antibody in a 1:500 dilution for 1 hour. Blots were washed again with TBST, developed with SuperSignal West Pico Chemiluminescent Substrate (Thermo Fisher Scientific), and imaged using the Biorad ChemiDoc MP Imaging System. Protein band densitometry was measured using the FIJI Plot Lanes function as area under the curve, and normalized to the respective internal control (β-Actin) band. Graphs and statistical analysis were performed in GraphPad prism.

Antibodies used for Western blot were as follows:

| Target | Host | Catalogue Number |
| --- | --- | --- |
| Phospho-Tau (Ser396) | Rabbit | Invitrogen 44-752G |
| Tau (4R isoform RD4) | Mouse | Sigma Aldrich 05-804 |
| Tau (HT7) | Mouse | Invitrogen MN1000 |
| β-Actin | Mouse | Sigma Aldrich A5316 |
| β-Actin | Rabbit | Sigma Aldrich ZRB1312 |
| HRP-linked anti-mouse IgG | Sheep | Cytiva NXA931 |
| HRP-linked anti-rabbit IgG | Donkey | Cytiva NA934 |

### Recording of action potentials and whole-cell currents

For electrophysiological recordings, miBrains were removed from the incubator and culture media was replaced with an artificial cerebrospinal fluid (ACSF) solution containing (in mM) 125 NaCl, 2.5KCl, 1.2 NaH2PO4•H2O, 2.4 CaCl2•2H2O, 1.2 MgCl2•6H2O, 26 NaHCO3 and 11 D-Glucose. miBrains were placed in a recording chamber and perfused with oxygenated ACSF at a constant rate of 2 mL/min at ∼32°C. Cells were visualized using infrared differential interference contrast imaging on an Olympus BX-50WI microscope.

Electrophysiological recordings were made using an Axon Multiclamp 700B patch-clamp amplifier (Molecular Devices) and Clampex software (version 11.2, Molecular Devices). Signals were filtered at 1 kHz using the amplifier’s four-pole, low-pass Bessel filter, digitized at 10 kHz with an Axon Digidata 1550B interface (Molecular Devices) and stored on a computer. Action potentials were generated by injecting various steps of currents in the whole-cell current clamp configuration and episodic stimulation acquisition mode. Whole-cell currents were recorded from a holding potential of −80 mV by stepping to various voltages using the whole-cell voltage clamp configuration and episodic stimulation acquisition mode. Recording electrodes pulled from borosilicate glass pipettes (World Precision Instruments) when filled with the internal solution (in mM): 120 K gluconate, 5 KCl, 2 MgCl2•6H2O, 10 HEPES, 4 ATP, 2 GTP. Data are presented as means ± standard errors of means.

## Acknowledgments

The authors would like to acknowledge the MIT Biology BioMicro Center for performing RNA-sequencing and quality analysis, the MIT Koch Institute Flow Cytometry Core at the Swanson Biotechnology Center for assistance with flow cytometry, and Matheus Victor for providing the mCherry iPSC line. The authors would also like to acknowledge generous support from The JPB Foundation, The Picower Institute for Learning and Memory, The BT Charitable Foundation, the Robert A. and Renee E. Belfer Family, Lester A. Gimpelson, Eduardo Eurnekian, the Halis Family, David B. Emmes, Kathleen and Miguel Octavio and, Jay L. And Carroll D. Miller.

## Funding

National Institutes of Health grant 3-UG3-NS115064-01S1 (LHT)

BT Charitable Foundation (LHT)

National Institutes of Health grant 1F32AG072813-01 (AES)

National Institutes of Health grant 1K01AG083734-01 (AES)

Kavanaugh Fellowship (AES)

Picower Fellowship (AB)

Schmidt Science Fellows in partnership with the Rhodes Trust (RLP)

## Author contributions

Conceptualization: LHT, JWB, AES, RL

Methodology: AES, AB, DSP, BJ

Investigation: AES, AB, EA, BJ, DSP, AJ, RLP, NT, CS

Visualization: AES, AB

Funding acquisition: LHT, JWB, MK, RL, AES

Project administration: LHT, RL

Supervision: LHT, RL

Writing – original draft: AES

Writing – review & editing: AES, LHT, RL, AB

## Data and materials availability

All data, code, and materials used in the analysis will be made available upon reasonable request.

