## Supporting Information for "Engineered 3D Immuno-Glial-Neurovascular Human miBrain Model"

**The file includes:**

Figs. S1 to S16
Captions for Movies S1 to S3

**Other Supplementary Materials for this manuscript include the following:**

Movies S1 to S3

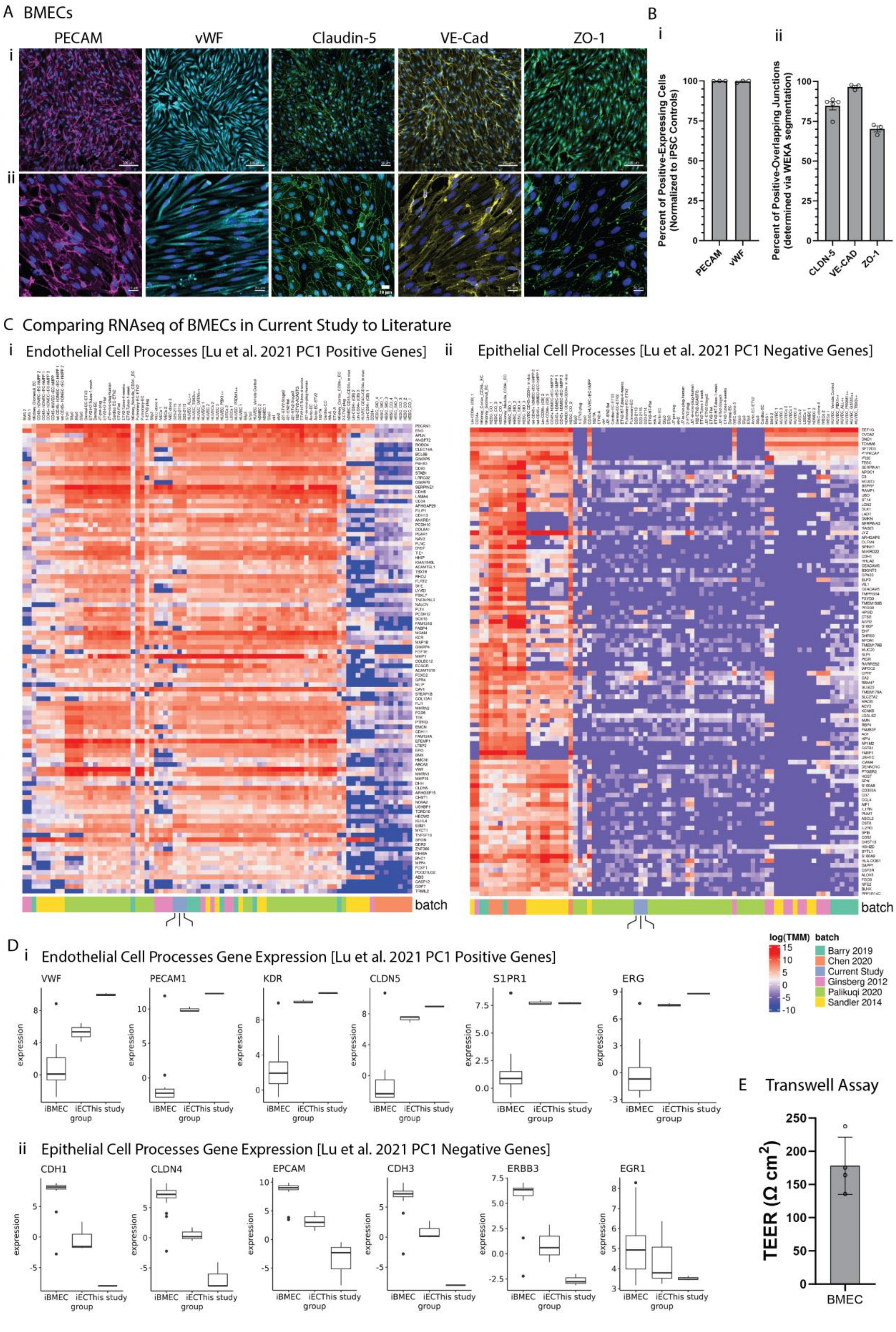
A BMECs
PECAM
vWF
Claudin-5
VE-Cad
ZO-1
i
ii
B
i
ii
Percent of Positive-Expressing Cells (Normalized to iPSC Controls)
PECAM
vWF
Percent of Positive-Overlapping Junctions (determined via WEKA segmentation)
CLDN-5
VE-CAD
ZO-1
C Comparing RNAseq of BMECs in Current Study to Literature
i Endothelial Cell Processes [Lu et al. 2021 PC1 Positive Genes]
ii Epithelial Cell Processes [Lu et al. 2021 PC1 Negative Genes]
batch
batch
log(TMM)
batch
Barry 2019
Chen 2020
Current Study
Ginsberg 2012
Palikuqi 2020
Sandler 2014
D i Endothelial Cell Processes Gene Expression [Lu et al. 2021 PC1 Positive Genes]
VWF
PECAM1
KDR
CLDN5
S1PR1
ERG
expression
iBMEC iECThis study
group
ii Epithelial Cell Processes Gene Expression [Lu et al. 2021 PC1 Negative Genes]
CDH1
CLDN4
EPCAM
CDH3
ERBB3
EGR1
E Transwell Assay
TEER (Ω cm²)
BMEC

**Fig. S1: Differentiation and Validation of BMECs.** (**A**) iPSC-derived BMECs immunoreactivity to endothelial cell markers (**i**) lower magnification images (*left to right*): PECAM (magenta: PECAM, blue: Hoechst; scale, 100 µm), vWF (cyan: vWF, blue: Hoechst; scale, 100 µm), Claudin-5 (green: Claudin-5, blue: Hoechst; scale, 50 µm), VE-Cad (yellow: VE-Cad, blue: Hoechst; scale bar, 100 µm), ZO-1 (green: ZO-1, blue: Hoechst; scale, 100 µm), (**ii**) higher magnification images as in *(i)* (scale bars, 20 µm), (**B**) quantification of the percent of positive marker-expressing cells via immunohistochemistry for (**i**) PECAM and vWF expression above iPSC controls and (**ii**) junctional markers Claudin-5, VE-Cad, and ZO-1 expression at borders defined by PECAM (Claudin-5) or phalloidin (VE-Cad, ZO-1) reference channel via WEKA segmentation and border overlay (n ≥ 3 wells per group), (**C**) BMEC RNAseq integrated to literature datasets for (**i**) endothelial cell processes and (**ii**) epithelial cell processes determined by Lu, et al. 2021 *(33)* with the sequencing from this study (batch legend: blue; indicated with lines below) integrated with literature datasets *(33-38)* (scale: blue to red for log(TMM) of -10 to 15), (**D**) gene expression for key genes from (**i**) endothelial cell processes and (**ii**) epithelial cell processes as analyzed in Lu, et al. 2021 *(33)* and comparing literature iBMECs and iECs to the BMECs used in this study, and (**E**) TEER assay for BMEC monolayers on 0.4 µm pore size, 12 mm diameter Transwell at day 15 (n = 4).

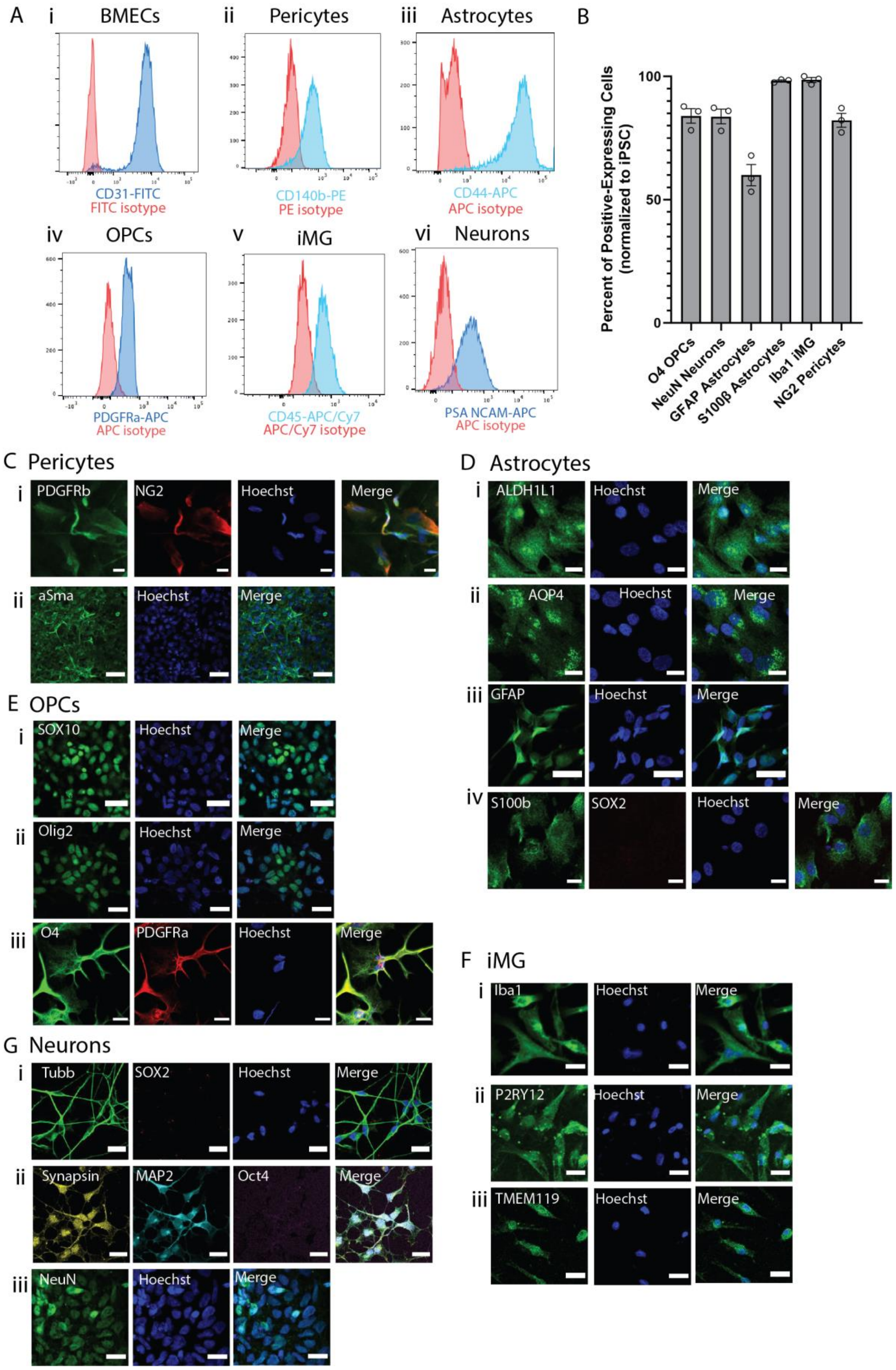
A
i BMECs
ii Pericytes
iii Astrocytes
CD31-FITC
FITC isotype
CD140b-PE
PE isotype
CD44-APC
APC isotype
iv OPCs
v iMG
vi Neurons
PDGFRa-APC
APC isotype
CD45-APC/Cy7
APC/Cy7 isotype
PSA NCAM-APC
APC isotype
B
Percent of Positive-Expressing Cells (normalized to iPSC)
100
50
0
O4 OPCs
NeuN Neurons
GFAP Astrocytes
S100β Astrocytes
Iba1 iMG
NG2 Pericytes
C Pericytes
i PDGFRb NG2 Hoechst Merge
ii aSma Hoechst Merge
D Astrocytes
i ALDH1L1 Hoechst Merge
ii AQP4 Hoechst Merge
iii GFAP Hoechst Merge
iv S100b SOX2 Hoechst Merge
E OPCs
i SOX10 Hoechst Merge
ii Olig2 Hoechst Merge
iii O4 PDGFRa Hoechst Merge
F iMG
i Iba1 Hoechst Merge
ii P2RY12 Hoechst Merge
iii TMEM119 Hoechst Merge
G Neurons
i Tubb SOX2 Hoechst Merge
ii Synapsin MAP2 Oct4 Merge
iii NeuN Hoechst Merge

**Fig. S2: Differentiation and Validation of iPSC-Derived Brain Cell Types.** (**A**) Flow cytometry marker analysis of (**i**) BMECs for canonical marker CD31 (blue: CD31, red: isotype control), (**ii**) iPSC-derived pericytes for CD140b (blue: CD140b, red: isotype control), (**iii**) iPSC-derived astrocytes for CD44 (blue: CD44, red: isotype control), (**iv**) iPSC-derived OPCs for PDGFRα (blue: PDGFRα, red: isotype control), (**v**) iPSC-derived iMG for CD45 (blue: CD45, red: isotype control), (**vi**) iPSC-derived neurons for excitatory cortical neuron marker PSA-NCAM (blue: PSA-NCAM, red: isotype control), (**B**) quantification of percent positive-expressing cells assessed via immunohistochemistry at the end of the differentiation protocols and two days after re-plating, compared to iPSCs stained in parallel, for key cell markers (n >= 3 wells/ marker), (**C**) representative immunofluorescence images of iPSC-derived pericytes expressing markers (**i**) PDGFRb and NG2 (green: PDGFRb, red: NG2, blue: Hoechst; scale bars, 20 µm) and (**ii**) αSMA (green: αSMA, blue: Hoechst; scale bars, 20 µm), (**D**) iPSC-derived astrocytes positive immunoreactivity to astrocyte markers (**i**) ALDH1L1 (green: ALDH1L1, blue: Hoechst; scale bar, 20 µm), (**ii**) AQP4 (green: AQP4, blue: Hoechst; scale bar, 20 µm), (**iii**) GFAP (green: GFAP, blue: Hoechst; scale bar, 20 µm), and (**iv**) S100b and absence of immunoreactivity to neuron progenitor marker SOX2 (green: S100b, red: SOX2, blue: Hoechst; scale bar, 20 µm), (**E**) iPSC-derived OPCs stained positive for markers (**i**) SOX10 (green: SOX10, blue: Hoechst; scale bar, 20 µm), (**ii**) Olig2 (green: Olig2, blue: Hoechst; scale bar, 20 µm), and (**iii**) O4 and PDGFRα (green: O4, red: PDGFRα, blue: Hoechst; scale bar, 20 µm), (**F**) iPSC-derived iMG immunoreactivity to canonical markers (**i**) Iba1 (green: Iba1, blue: Hoechst; scale bar, 20 µm), (**ii**) P2RY12 (green: P2RY12, blue: Hoechst; scale bar, 20 µm), and (**iii**) TMEM119 (green: TMEM119, blue: Hoechst; scale bar, 20 µm), and (**G**) iPSC-derived neurons immunoreactivity (**i**) to b-Tubulin and SOX2 (green: TUBB, red: SOX2, blue: Hoechst; scale bar, 20 µm), (**ii**) MAP2 and synapsin and absence of immunoreactivity to pluripotency marker OCT4 (yellow: synapsin, cyan: MAP2, magenta: OCT4, blue: Hoechst; scale bars, 20 µm), and (**iii**) NeuN (green: NeuN, blue: Hoechst; scale bar, 20 µm).

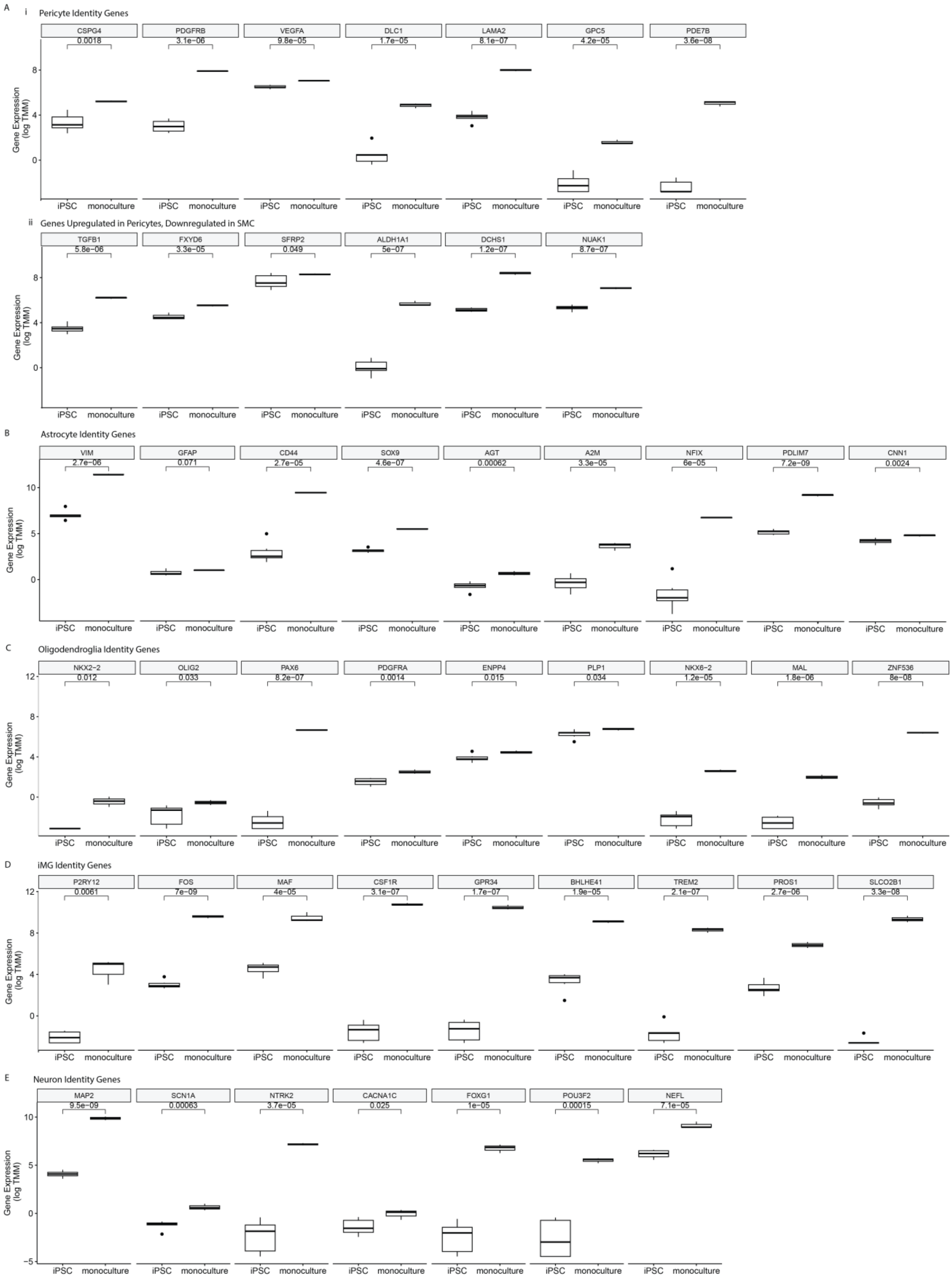
A
i Pericyte Identity Genes
CSPG4
PDGFRB
VEGFA
DLC1
LAMA2
GPC5
PDE7B
0.0018
3.1e−06
9.8e−05
1.7e−05
8.1e−07
4.2e−05
3.6e−08
Gene Expression (log TMM)
iPSC
monoculture
ii Genes Upregulated in Pericytes, Downregulated in SMC
TGFB1
FXYD6
SFRP2
ALDH1A1
DCHS1
NUAK1
5.8e−06
3.3e−05
0.049
5e−07
1.2e−07
8.7e−07
B
Astrocyte Identity Genes
VIM
GFAP
CD44
SOX9
AGT
A2M
NFIX
PDLIM7
CNN1
2.7e−06
0.071
2.7e−05
4.6e−07
0.00062
3.3e−05
6e−05
7.2e−09
0.0024
C
Oligodendroglia Identity Genes
NKX2−2
OLIG2
PAX6
PDGFRA
ENPP4
PLP1
NKX6−2
MAL
ZNF536
0.012
0.033
8.2e−07
0.0014
0.015
0.034
1.2e−05
1.8e−06
8e−08
D
iMG Identity Genes
P2RY12
FOS
MAF
CSF1R
GPR34
BHLHE41
TREM2
PROS1
SLCO2B1
0.0061
7e−09
4e−05
3.1e−07
1.7e−07
1.9e−05
2.1e−07
2.7e−06
3.3e−08
E
Neuron Identity Genes
MAP2
SCN1A
NTRK2
CACNA1C
FOXG1
POU3F2
NEFL
9.5e−09
0.00063
3.7e−05
0.025
1e−05
0.00015
7.1e−05

**Fig. S3: Gene Expression Validation of iPSC-Derived Brain Cell Types.** (**A**) Gene expression for iPSCs and pericyte monocultures for (**i**) key pericyte identity genes and (**ii**) genes that have been found to be upregulated in pericytes and downregulated in SMCs, (**B**) gene expression for iPSCs and astrocyte monocultures for astrocyte identity genes, (**C**) gene expression for iPSCs and OPC monocultures for OPC and oligodendrocyte identity genes, (**D**) gene expression for iPSCs and iMG monocultures for microglial identity genes, and (**E**) gene expression for iPSCs and neuron monocultures for neuron identity genes (expression plotted as log TMM; statistical significance determined via t test).

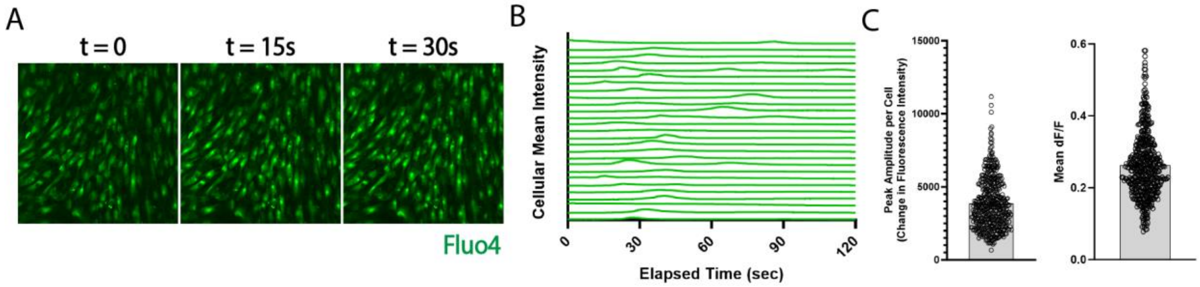


**Fig. S4: Characterization of Astrocyte Calcium Transients.** (**A**) Images of calcium transients recorded from astrocyte monocultures with calcium indicator Fluo4, (**B**) representative traces of calcium transients plotting mean intensity over time, and (**C**) quantification of (left) peak amplitude per cell in terms of fluorescence intensity and (right) mean dF/F for n = 485 cells.

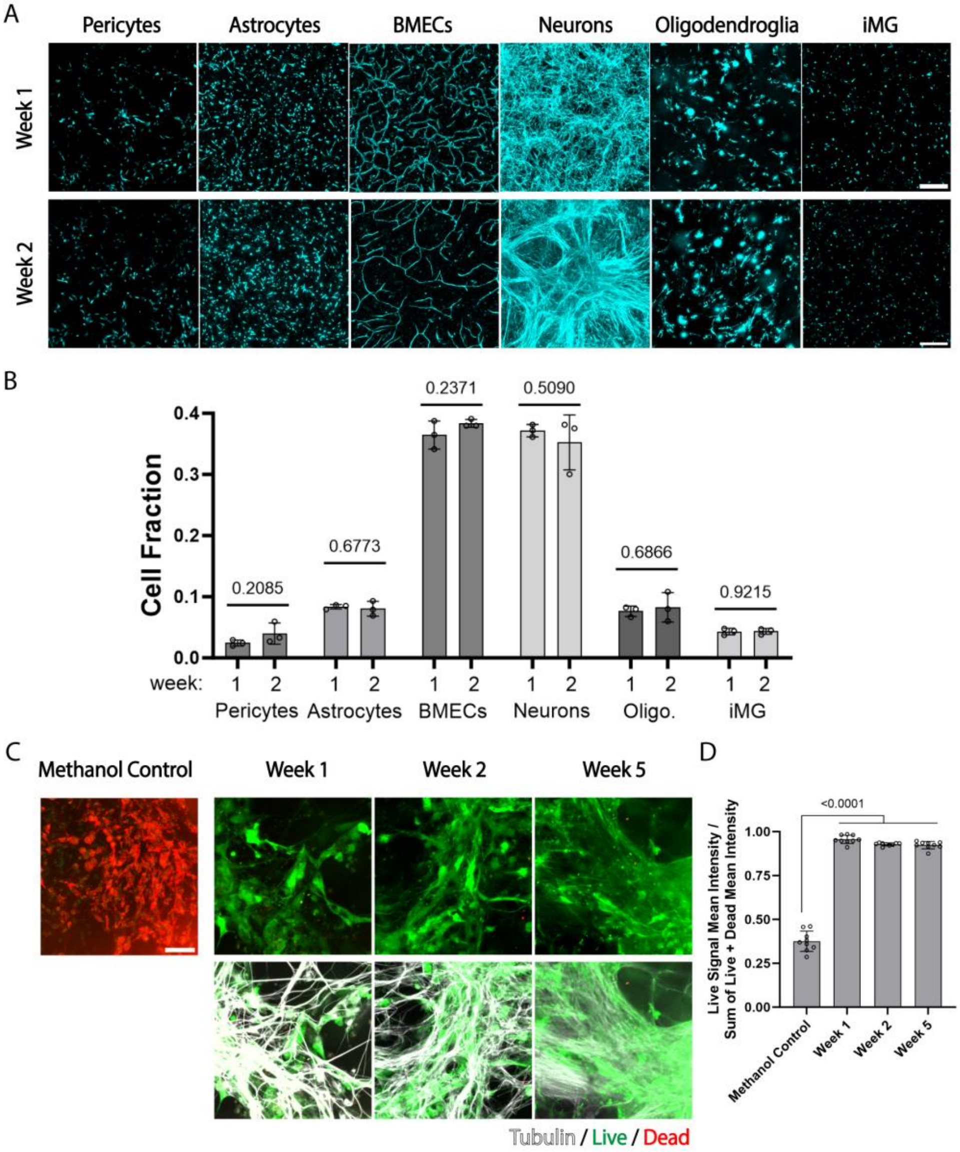


**Fig. S5: miBrain Cell Composition.** (**A**) miBrain cell types (*left to right*) pericytes (cyan: mCherry-pericytes), astrocytes (cyan: mCherry-astrocytes), BMECs (cyan: mCherry-BMECs), neurons (cyan: tubulin), oligodendroglia (cyan: tdTomato-oligodendroglia), and iMG (cyan: membrane-labeled iMG) for (*top*) week 1 and (*bottom*) week 2, (**B**) quantification of cell fractions of cell types at weeks 1 and 2 (n = 3 miBrains/ cell type/ group, repeated in 3 independent batches; one-way ANOVA statistical test with multiple comparisons), (**C**) live/dead assay for miBrains at weeks 1, 2, and 5 (*top*) compared to methanol control (green: live, red: dead; scale bar, 100 µm) and (*bottom*) co-labeled with tubulin live dye to visualize neurons (gray: tubulin, green: live, red: dead), and (**D**) live cell fraction in each condition (n = 3 fields of view from n = 3 miBrains/ group, repeated in 3 independent batches; statistical significance determined via t test).

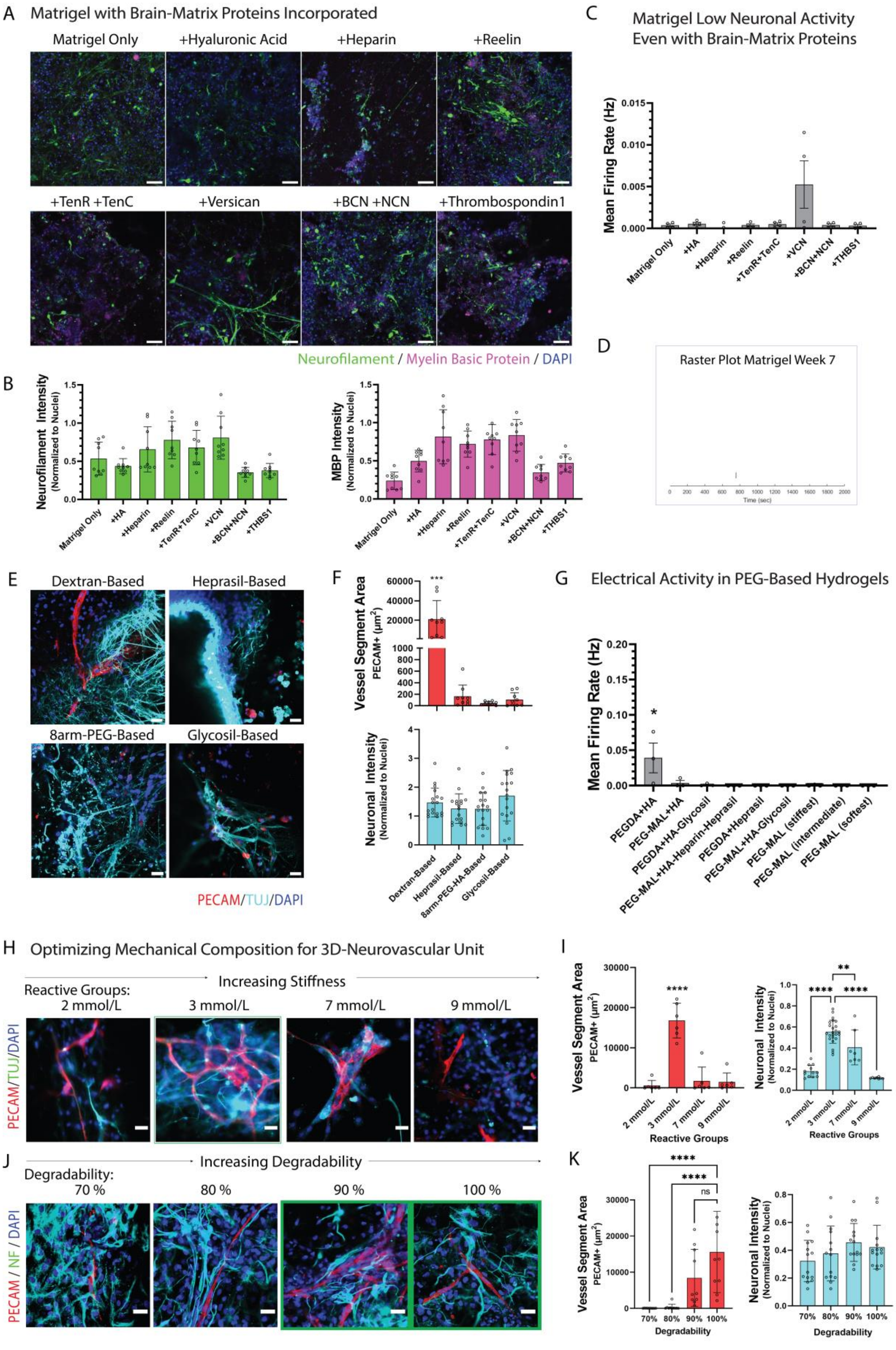
A
Matrigel with Brain-Matrix Proteins Incorporated
Matrigel Only
+Hyaluronic Acid
+Heparin
+Reelin
+TenR +TenC
+Versican
+BCN +NCN
+Thrombospondin1
Neurofilament / Myelin Basic Protein / DAPI
B
Neurofilament Intensity (Normalized to Nuclei)
MBP Intensity (Normalized to Nuclei)
Matrigel Only
+HA
+Heparin
+Reelin
+TenR+TenC
+VCN
+BCN+NCN
+THBS1
C
Matrigel Low Neuronal Activity Even with Brain-Matrix Proteins
Mean Firing Rate (Hz)
D
Raster Plot Matrigel Week 7
Time (sec)
E
Dextran-Based
Heprasil-Based
8arm-PEG-Based
Glycosil-Based
PECAM/TUJ/DAPI
F
Vessel Segment Area PECAM+ (μm²)
Neuronal Intensity (Normalized to Nuclei)
Dextran-Based
Heprasil-Based
8arm-PEG-HA-Based
Glycosil-Based
G
Electrical Activity in PEG-Based Hydrogels
Mean Firing Rate (Hz)
PEGDA+HA
PEG-MAL+HA
PEGDA+HA-Glycosil
PEG-MAL+HA-Heparin-Heprasil
PEGDA+Heprasil
PEG-MAL+HA-Glycosil
PEG-MAL (stiffest)
PEG-MAL (intermediate)
PEG-MAL (softest)
H
Optimizing Mechanical Composition for 3D-Neurovascular Unit
Increasing Stiffness
Reactive Groups:
2 mmol/L
3 mmol/L
7 mmol/L
9 mmol/L
PECAM/TUJ/DAPI
I
Vessel Segment Area PECAM+ (μm²)
Neuronal Intensity (Normalized to Nuclei)
Reactive Groups
J
Increasing Degradability
Degradability:
70 %
80 %
90 %
100 %
PECAM / NF / DAPI
K
Vessel Segment Area PECAM+ (μm²)
Neuronal Intensity (Normalized to Nuclei)
Degradability

**Fig. S6: Engineering a Brain-Mimicking Hydrogel for Multicellular miBrain Co-Culture.** (**A**) Neuronal morphology (green: neurofilament) and myelination (magenta: MBP) in miBrains encapsulated in Matrigel incorporated with various brain extracellular matrix proteins (scale bars, 50 µm), (**B**) average intensity of neuronal (top) and MBP immunoreactivity (bottom) across miBrains encapsulated in Matrigel (data from n = 3 replicates, n = 3 fields of view each; plotted as mean and S.D.), (**C**) neuronal activity as measured on an MEA at week 7 for mean firing rate (data from n = 4 replicates, averaged over recordings of at least 30 min.; plotted as mean and S.E.M.), (**D**) raster plot for spikes of neuronal activity in a representative miBrain in Matrigel at week 7, (**E**) single z-plane images of neuronal (cyan: Tuj1) and vessel morphology (red: PECAM) after 2 weeks in culture in various engineered hydrogels (scale bars, 30 µm), (**F**) quantified for area of the segments of vessels (PECAM-positive) captured in images (top; data from n = 3 replicates, n = 3 fields of view each; plotted as mean and S.D.; statistical analysis via one-way ANOVA, *** $p = 0.003$) and the Tuj1 immunoreactivity (bottom; data from n = 3 replicates, n = 6 fields of view each), (**G**) neuronal activity of miBrains encapsulated in various engineered hydrogels, as assessed by an MEA system after 2 weeks in culture (data from n = 3 replicates, averaged over recordings of at least 30 min.; plotted as mean and S.E.M.; statistical analysis via one-way ANOVA, * $p<0.05$), (**H**) optimization of mechanical properties of dextran-based hydrogels across amounts of macromer and crosslinker reactive groups, screening across 3D, integral neurovascular network co-assembly for vessels (red: PECAM) and neurons (cyan: Tuj1) with single z-plane images (scale bars, 30 µm), (**I**) quantified in terms of area of the segments of vessels (PECAM+) captured in images (left; data from n = 3 replicates, n = 6 field of view, mean and S.D.; statistical analysis via one-way ANOVA, **** $p < 0.0001$) and the Tuj1 immunoreactivity (right; data from n = 3 replicates, n = 7 fields of view; statistical analysis via one-way ANOVA, mean and S.D., ** $p = 0.006$, **** $p < 0.0001$), (**J**) varying degradability in dextran-based hydrogels, screening across 3D, integral neurovascular network co-assembly for vessels (red: PECAM) and neurons (cyan: Tuj1) with single z-plane images (scale bars, 30 µm), (**K**) quantified in terms of area of the segments of vessels (PECAM+) captured in images (left; data from n = 3 replicates, n = 10 fields of view, mean and S.D.; statistical analysis via one-way ANOVA, **** $p < 0.0001$) and the Tuj1 immunoreactivity (right; data from n = 3 replicates, n = 3 fields of view each, mean and S.D.).

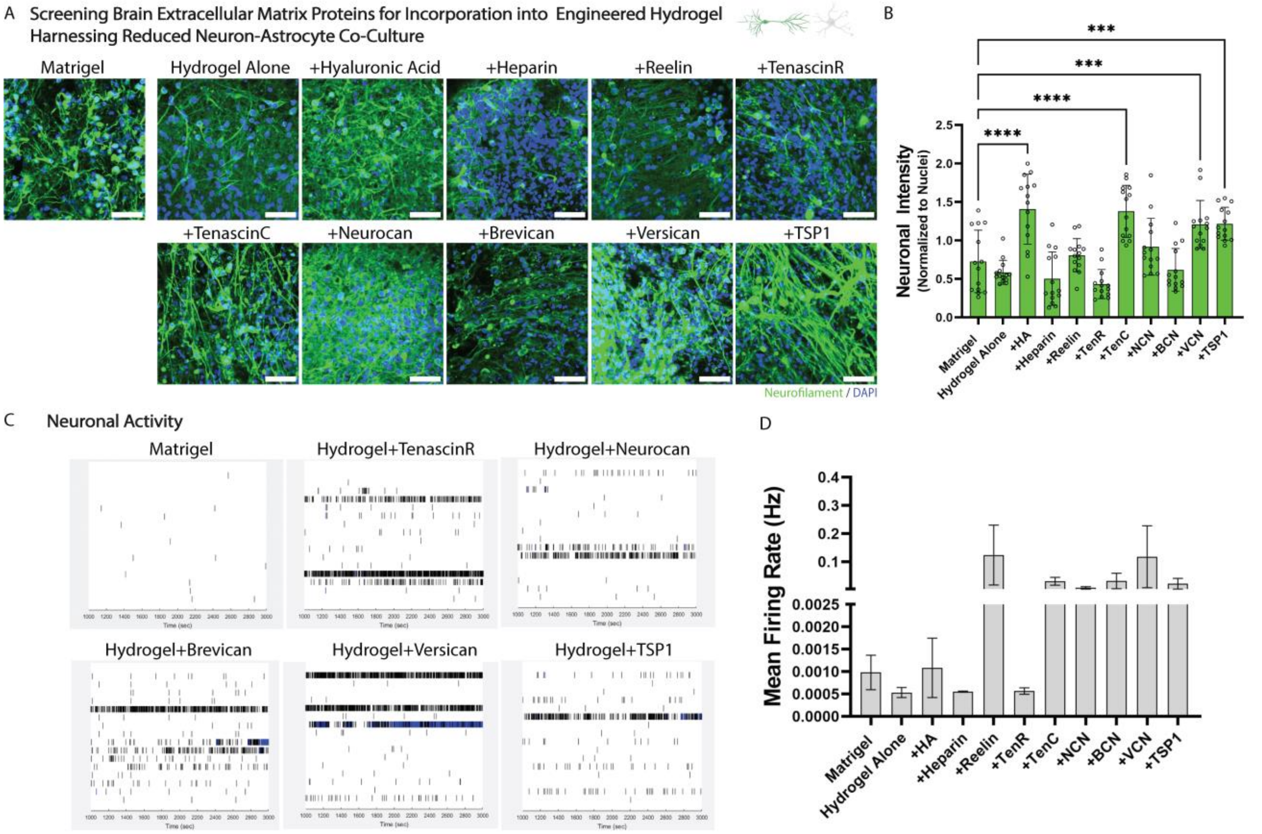


**Fig. S7: Brain Extracellular Matrix Components Incorporated into Engineered Hydrogel to Enhance Neuronal Phenotypes.** (**A**) Neuronal morphology (green: neurofilament) after 2 weeks in a neuron-astrocyte co-culture used to screen across dextran-based engineered hydrogels incorporated with various brain extracellular matrix proteins (scale bars, 50 µm), (**B**) average intensity of neuronal immunoreactivity across conditions (data from n = 3 replicates, n = 14 fields of view; statistical analysis via one-way ANOVA, *** p = 0.006, **** p < 0.0001), (**C**) example raster plots of neuronal activity assessed on an MEA system early in the culture at week 2, and (**D**) averaged mean firing rate at week 2 across extracellular matrix conditions (data are from n = 3 replicates, averaged over recordings of at least 30 min.; mean and S.E.M.).

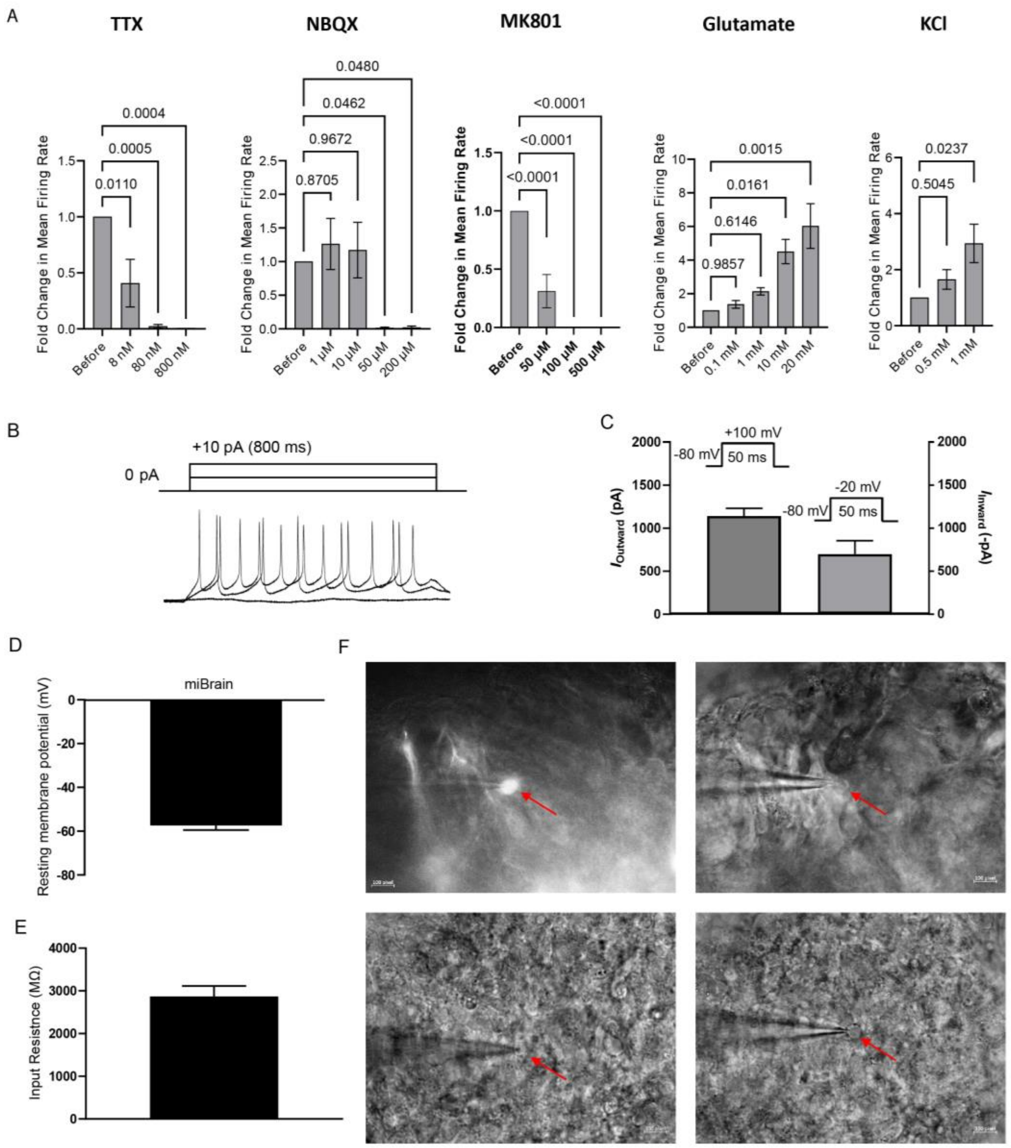


**Fig. S8: Neuromodulator and Electrophysiological Analysis of miBrain Neurons.** (**A**) Change in mean firing rate for miBrains in response to varied doses of neuromodulators assessed via MEA for (*from left to right*) TTX, NBQX, MK801, MNI-caged-L-Glutamate, and KCl (n = 4 miBrains/ group; plotted as mean and S.E.M.; statistical significance determined via one-way ANOVA), (**B**) representative whole-cell current clamp recording of action potentials from miBrain-neurons, (**C**) summary of whole-cell voltage clamp recording of inward and outward currents (n = 31 cells), (**D**) summary of resting membrane potentials (n = 31 cells), (**E**) summary of input resistance (n = 31 cells), and (**F**) images of whole-cell patch clamp recording of miBrain-neurons.

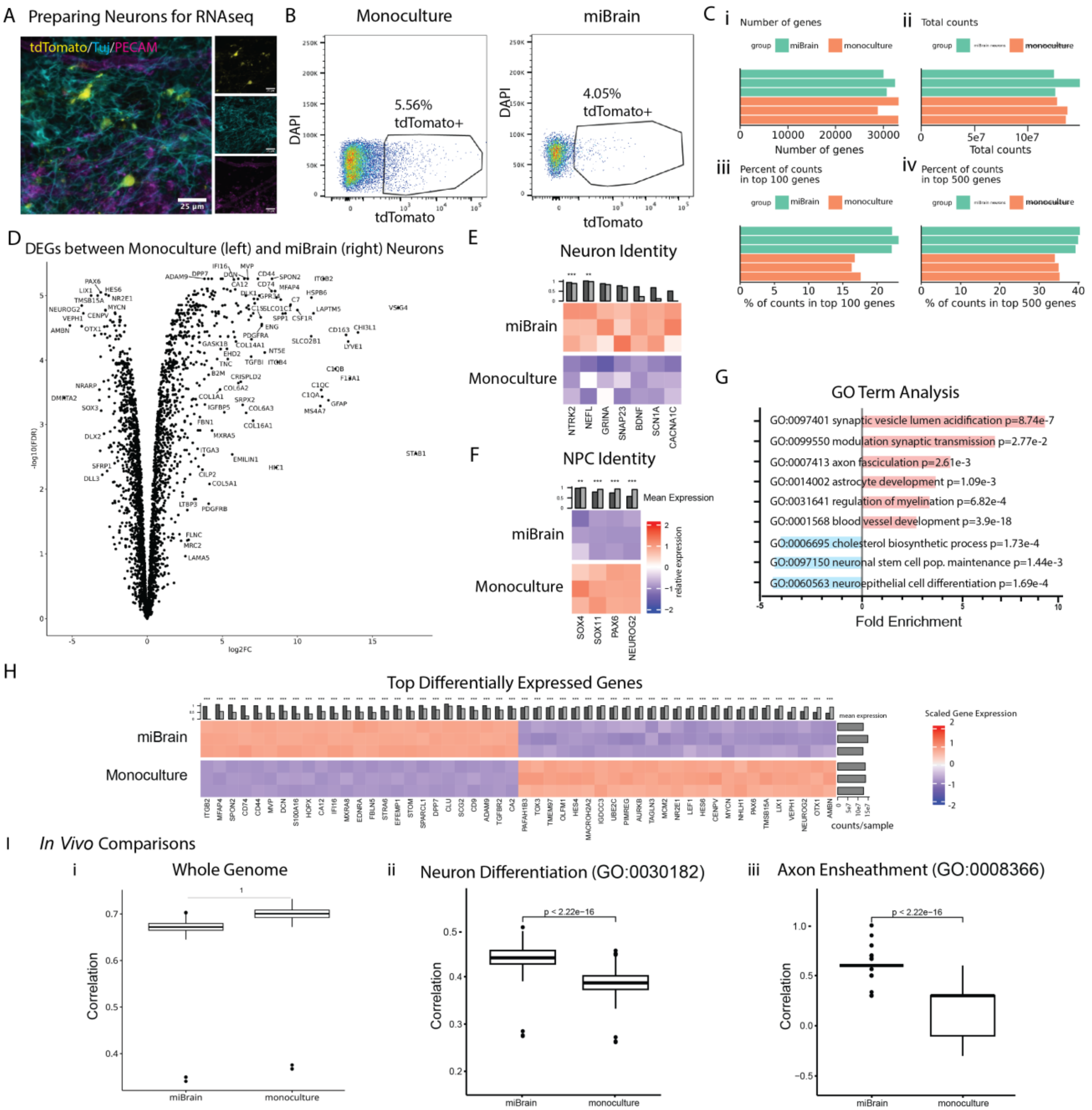


**Fig. S9: Transcriptomic Analysis of miBrain-Neurons.** (**A**) tdTomato-neurons incorporated into miBrains enabling tdTomato-based isolation of neurons via flow cytometry (yellow: tdTomato, magenta: PECAM, cyan: Tuj; scale bars, 25 μm), (**B**) gating of tdTomato-positive cells in flow cytometry (**C**) RNA-sequencing of neurons in miBrains and neurons monocultures: (**i**) genes per sample, (**ii**) total gene counts, (**iii**) percent of gene counts in the top 100 genes, and (**iv**) percent of gene counts in the top 500 genes, (**D**) upregulation in the miBrain-neurons of genes associated with neuron identity, (**E**) downregulation upregulation in the miBrain-neurons of genes associated with NPC identity, (**F**) top DEGs between neurons isolated from miBrains versus monocultures sorted by mean expression in miBrain, plotting TMM-normalized and scaled expression (FDR < 0.001), (**G**) volcano plot displaying these top DEGs between monocultured and miBrain neurons (left: upregulated in monocultured neurons, right: upregulated in miBrain- neurons), and (**H**) gene ontology analysis of iMG RNAseq for biological pathways significantly altered in miBrain-cultured iMG based on significantly upregulated and downregulated DEGs and with an FDR p-value less than 0.05, and (**I**) correlation of miBrain-

and monoculture-neurons to human *in vivo* prefrontal cortex excitatory neurons identified via snRNAseq of non-AD decedent tissue *(47)* for the (**i**) whole genome, (**ii**) genes associated with neuron differentiation (GO:0030182), and (**iii**) genes associated with axon ensheathment (GO:0008366) (statistical significance determined via t test).

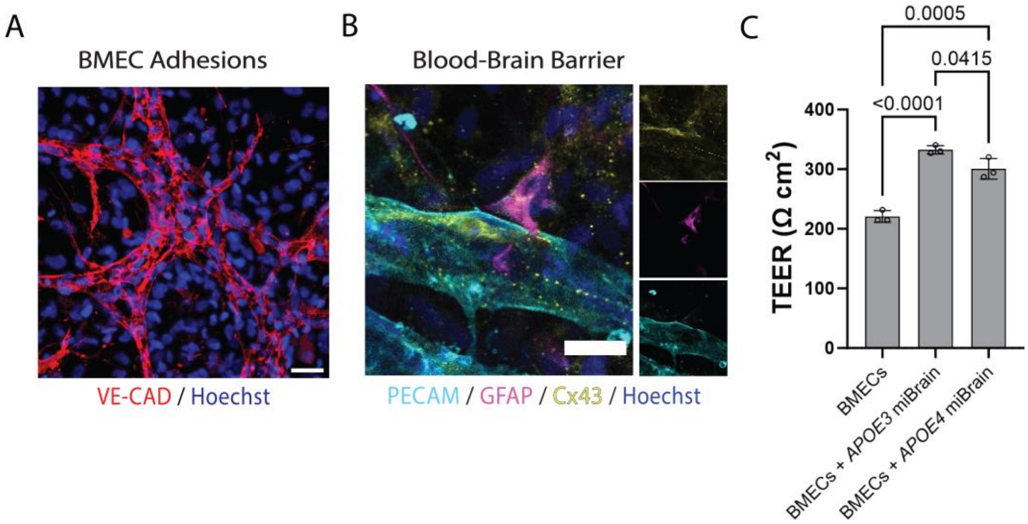


**Fig. S10: Marker and Functional BBB Characterization.** (**A**) VE-CAD adhesions at vessels (red: VE-CAD, blue: Hoechst; scale bar, 30 µm), (**B**) astrocytes at microvasculature and BBB receptor immunohistochemistry at vessels (yellow: Cx43, cyan: PECAM, magenta: GFAP, blue: Hoechst; scale bar, 25 µm), and (**C**) characterization of TEER for BMEC monocultures versus BMEC monocultures seeded with *APOE3* miBrains versus BMEC monocultures seeded with *APOE4* miBrains, measured 6 days following miBrain seeding (day 10 in culture overall; n = 3/ group; statistical significance determined via one-way ANOVA).

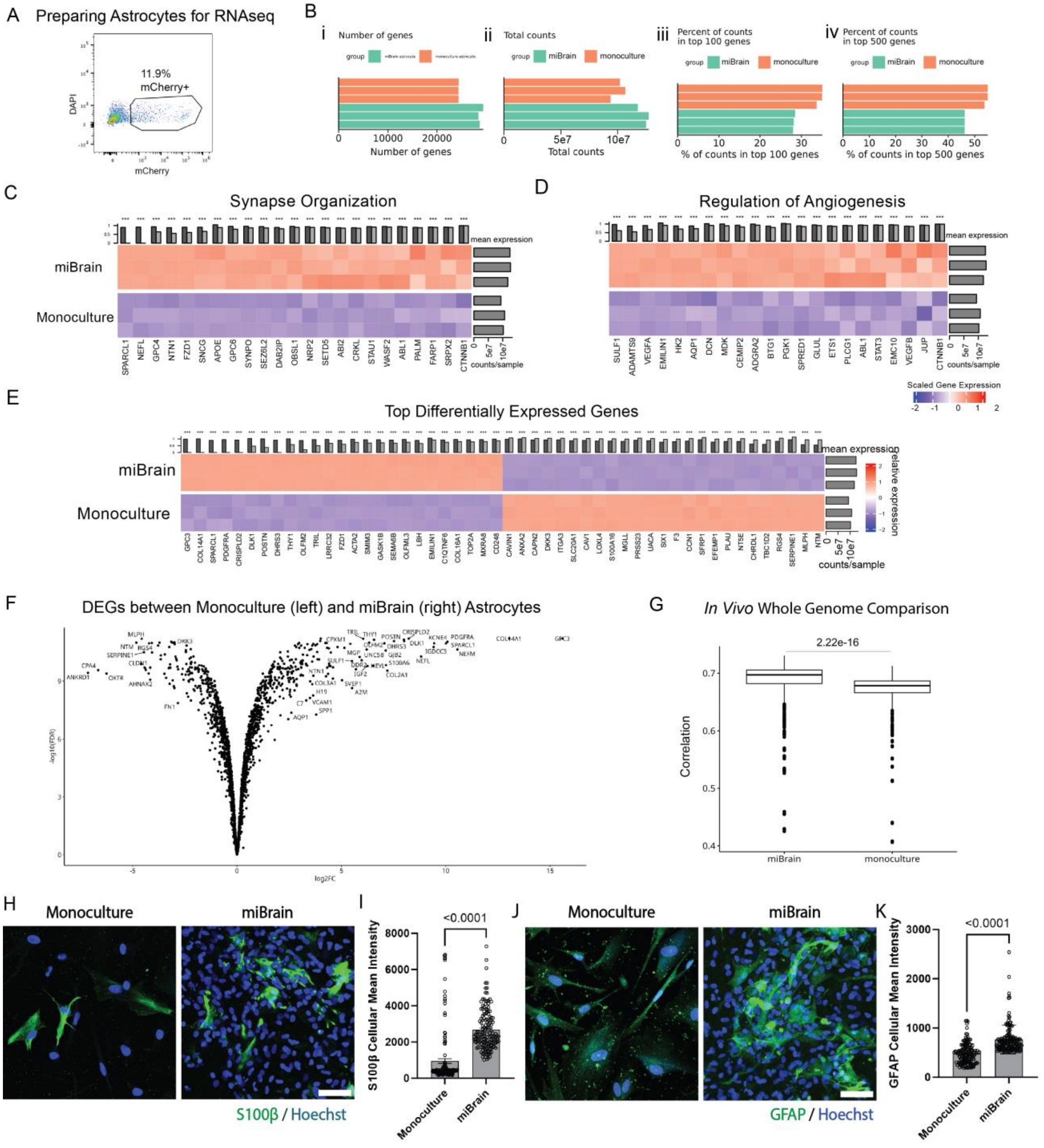


**Fig. S11: Transcriptomic Analysis of miBrain-Astrocytes.** (**A**) Gating of mCherry-positive cells in flow cytometry to identify astrocytes, (**B**) RNA-sequencing of astrocytes in miBrains and astrocyte monocultures: (**i**) genes per sample, (**ii**) total gene counts, (**iii**) percent of gene counts in the top 100 genes, and (**iv**) percent of gene counts in the top 500 genes, (**C**) upregulation in the miBrain-astrocytes of genes associated with synapse organization (GO:0050808; * $p < 0.05$, ** $p < 0.01$, *** $p < 0.001$), (**D**) upregulation in the miBrain-astrocytes of genes associated with regulation of angiogenesis (GO:0045765; * $p < 0.05$, ** $p < 0.01$, *** $p < 0.001$), (**E**) top DEGs between astrocytes isolated from miBrains versus monocultures sorted by mean expression in miBrain, plotting TMM-normalized and scaled expression (FDR < 0.001), and (**F**) volcano plot displaying these top DEGs between monocultured and miBrain astrocytes (left: upregulated in monocultured astrocytes, right: upregulated in miBrain-astrocytes), (**G**) correlation of miBrain- and monoculture-astrocytes to human *in vivo* prefrontal cortex astrocytes identified via snRNAseq of non-AD decedent tissue referencing the whole genome *(47)* (statistical significance determined via t test), (**H**) immunoreactivity to S100b in astrocyte monocultures compared to miBrains (green: S100b, blue: Hoechst; scale bar, 50 µm), (**I**) quantification of mean intensity of S100b-positive cells (n = 150 cells across n = 3 wells/ group; reported as mean and S.E.M.; statistical significance determined via t test), (**J**) immunoreactivity to GFAP in astrocyte monocultures compared to miBrains (green: GFAP, blue: Hoechst; scale bar, 50 µm), and (**K**) quantification of mean intensity of GFAP-positive cells (n = 150 cells across n = 3 wells/ group; reported as mean and standard deviation; statistical significance determined via t test).

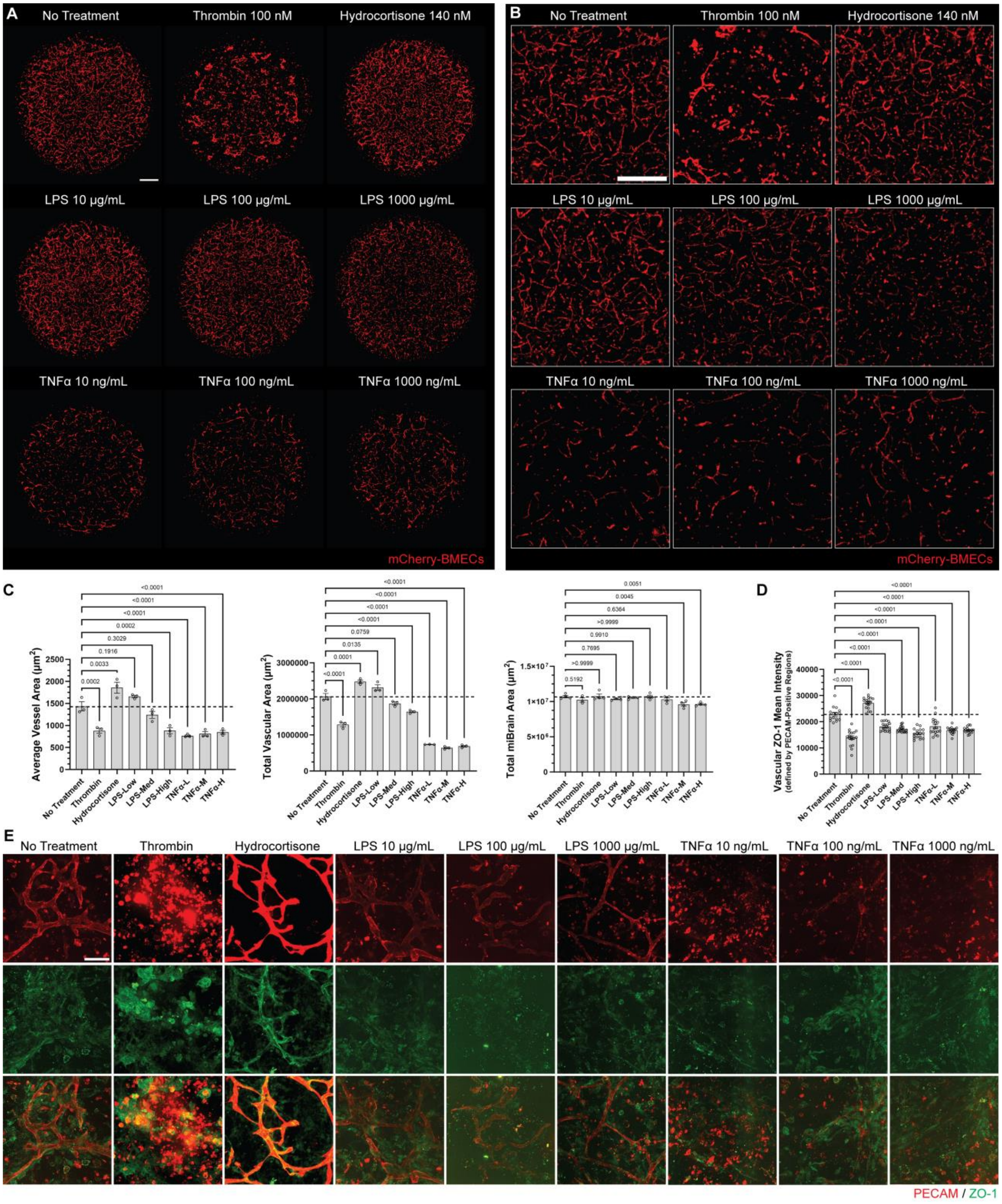


**Fig. S12: Functional Barrier Assessment with BBB Modulators.** (**A**) Whole-miBrain images and (**B**) magnified views of miBrain vasculature following 24 hour treatment with 100 nM thrombin, 140 nM hydrocortisone, 10, 100, or 1000 µg/mL LPS, and 10, 100, or 1000 ng/mL TNFα (red: mCherry-BMECs; scale bars, 500 µm), (**C**) quantification of (left) average vessel area, (middle) total vascular area, and (right) total miBrain area (n = 3/ group; statistical significance determined via one-way ANOVA), (**D**) quantification of ZO-1 mean signal intensity within vessel regions across treatment groups at 48 hours (n >= 15 ROIs/ group; statistical significance determined via one-way ANOVA), and (**E**) corresponding representative images across treatment groups at 48 hours (red: PECAM, green: ZO-1; scale bar, 200 µm).

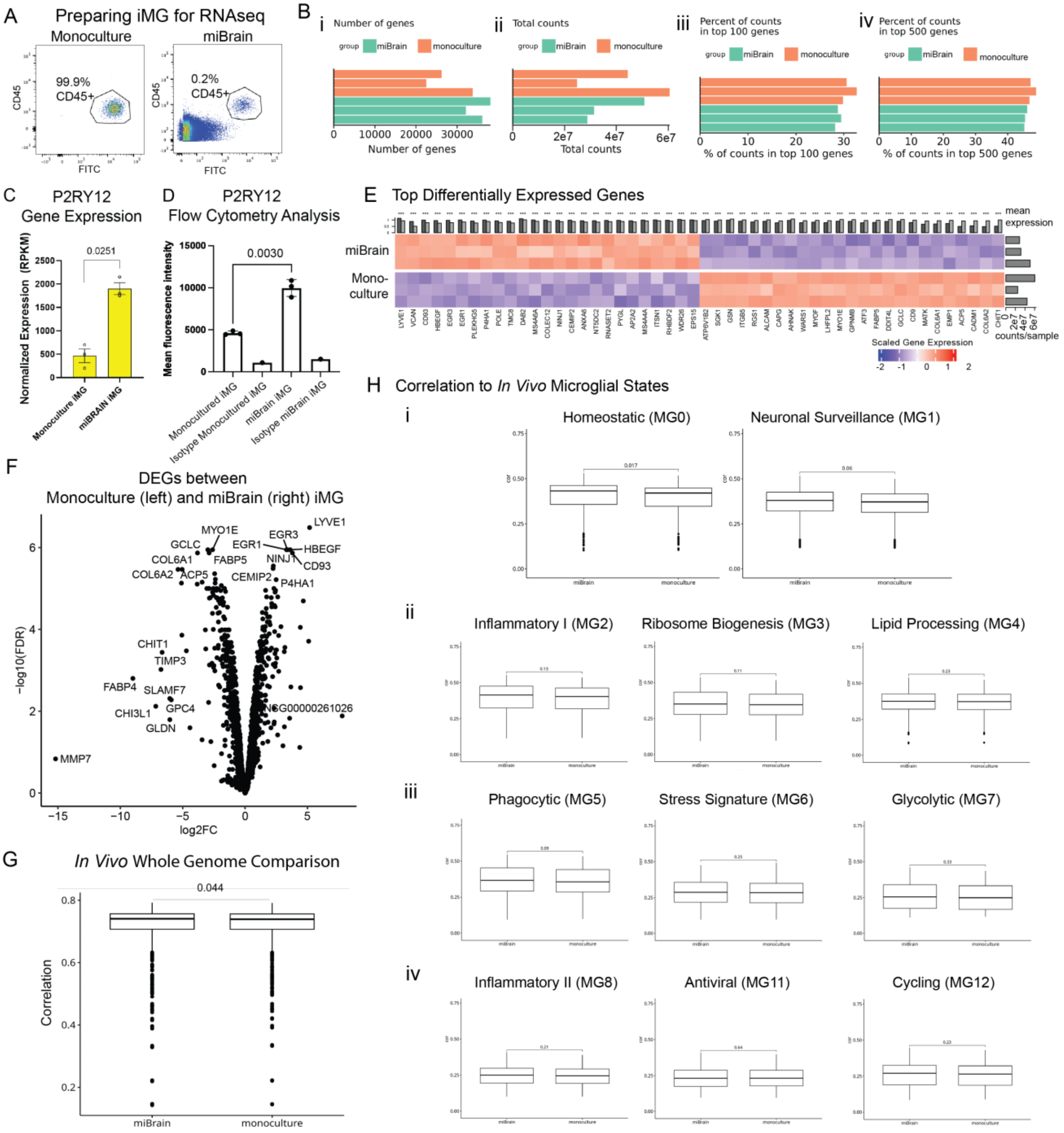


**Fig. S13: Transcriptomic Analysis of miBrain-iMG.** (**A**) iMG were isolated from miBrains using flow cytometry-based sorting of CD45-positive cells, (**B**) RNA-sequencing of iMG in miBrains and iMG monocultures: (**i**) genes per sample, (**ii**) total gene counts, (**iii**) percent of gene counts in the top 100 genes, and (**iv**) percent of gene counts in the top 500 genes, (**C**) gene expression of *P2RY12* in iMG for monocultures compared to miBrain, and (**D**) validation of P2RY12 differences in iMG between monoculture and miBrain via FACS analysis at the protein level, (**E**) top DEGs between iMG isolated from miBrains versus monocultures sorted by mean expression in miBrain, plotting TMM-normalized and scaled expression (FDR < 0.001), (**F**) volcano plot displaying these top DEGs between monocultured and miBrain iMG (left: upregulated in monocultured iMG, right: upregulated in miBrain-iMG), (**G**) correlation of miBrain- and monoculture-iMG to human *in vivo* prefrontal cortex microglia identified via snRNAseq of non-AD decedent tissue referencing the whole genome *(47)* (statistical significance determined via t test), and (**H**) correlation of miBrain- and monoculture-iMG to various microglial states for homeostatic and AD-pathological processes determined via snRNAseq across brain regions *(48)* (**i**) homeostatic and neuronal surveillance, (**ii**) inflammatory I, ribosome biogenesis, lipid processing, (**iii**) phagocytic, stress signature, glycolytic, (**iv**) inflammatory II, antiviral, and cycling states (statistical significance determined via t test).

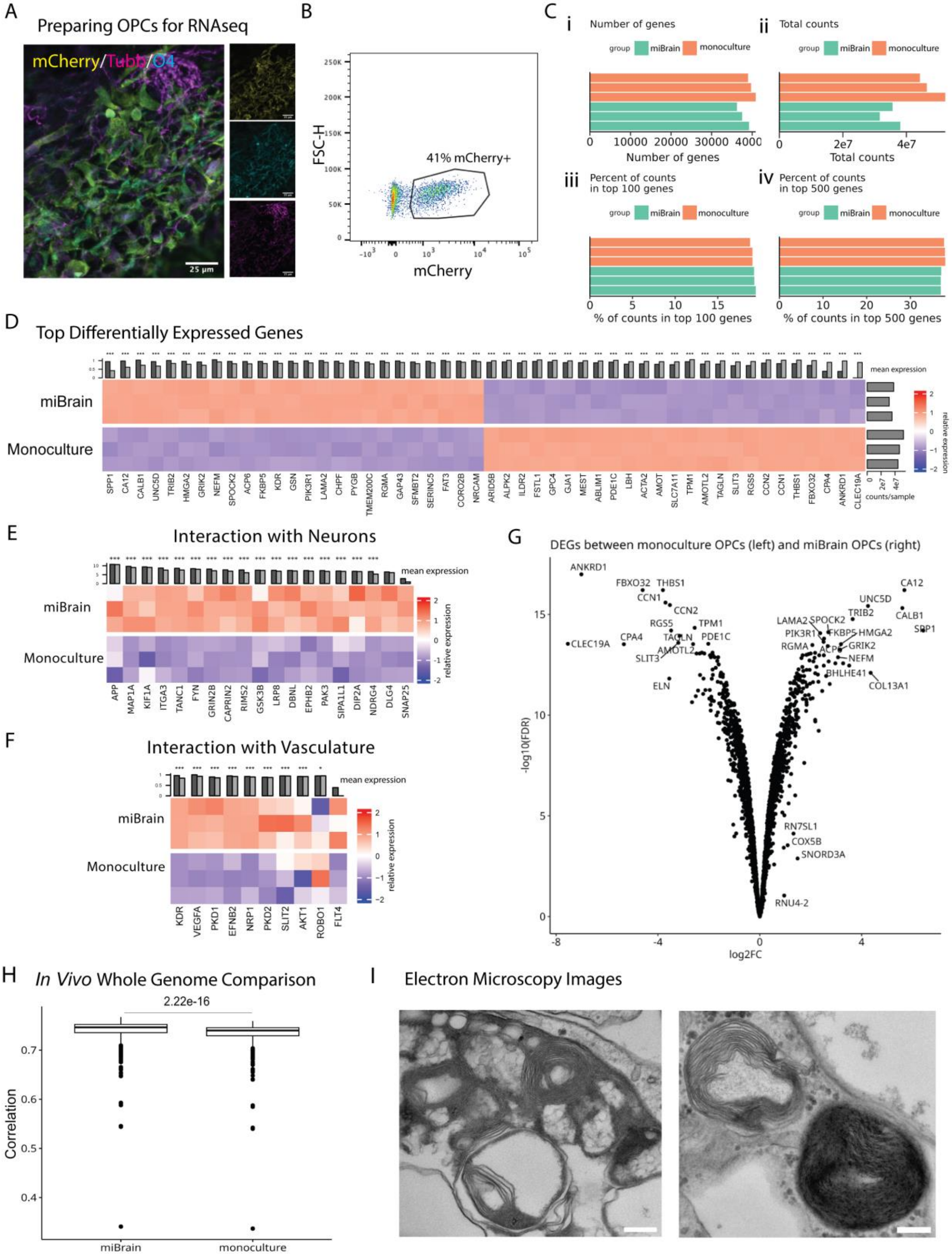


**Fig. S14: Transcriptomic Analysis of miBrain-Oligodendroglia.** (**A**) mCherry-oligodendroglia incorporated into miBrains enabling mCherry-based isolation of oligodendroglia via flow cytometry (yellow: mCherry, magenta: TUBB, cyan: O4; scale bars, 25 µm), (**B**) gating of mCherry-positive cells in flow cytometry (**C**) RNA-sequencing of oligodendroglia in miBrains and oligodendroglia monocultures: (**i**) genes per sample, (**ii**) total gene counts, (**iii**) percent of gene counts in the top 100 genes, and (**iv**) percent of gene counts in the top 500 genes, (**D**) top DEGs between oligodendroglia isolated from miBrains versus monocultures sorted by mean expression in miBrain, plotting TMM-normalized and scaled expression (FDR < 0.001), (**E**) expression of key *in vivo*-like genes in RNAseq of oligodendroglia isolated from miBrain compared to monocultured oligodendroglia displayed as TMM-normalized and scaled expression for interactions with neurons and (**F**) interactions with vasculature, (**G**) volcano plot displaying these top DEGs between monocultured and miBrain oligodendroglia (left: upregulated in monocultured oligodendroglia, right: upregulated in miBrain-oligodendroglia), and (**H**) correlation of miBrain- and monoculture-oligodendroglia to human *in vivo* prefrontal cortex OPCs identified via snRNAseq of non-AD decedent tissue referencing the whole genome *(47)* (statistical significance determined via t test), and (**I**) electron microscopy images of miBrain (scale bar, 200 µm).

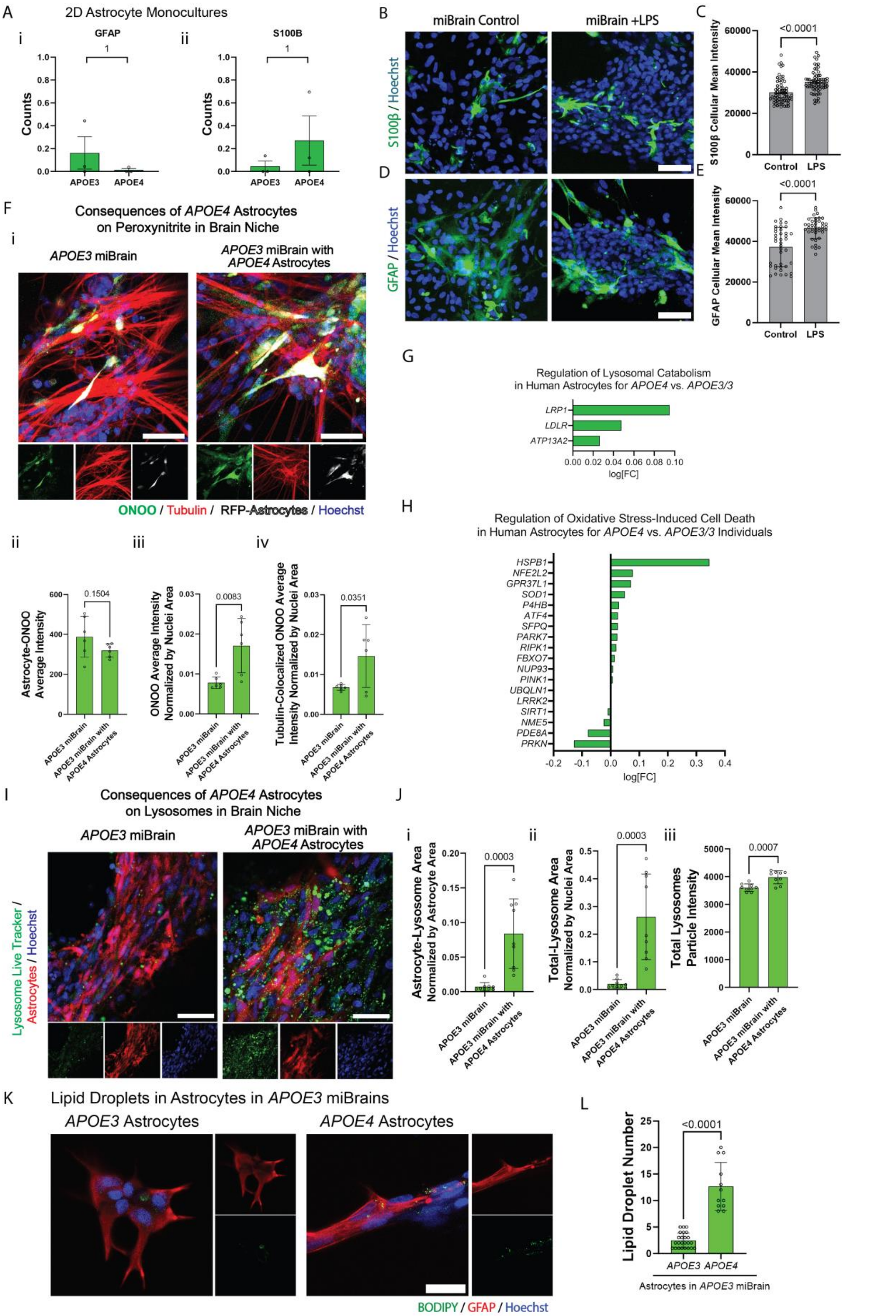
A
2D Astrocyte Monocultures
i
GFAP
ii
S100B
Counts
APOE3
APOE4
B
miBrain Control
miBrain +LPS
S100β / Hoechst
C
S100β Cellular Mean Intensity
<0.0001
Control
LPS
D
GFAP / Hoechst
E
GFAP Cellular Mean Intensity
<0.0001
F
Consequences of APOE4 Astrocytes on Peroxynitrite in Brain Niche
APOE3 miBrain
APOE3 miBrain with APOE4 Astrocytes
ONOO / Tubulin / RFP-Astrocytes / Hoechst
ii
iii
iv
Astrocyte-ONOO Average Intensity
0.1504
ONOO Average Intensity Normalized by Nuclei Area
0.0083
Tubulin-Colocalized ONOO Average Intensity Normalized by Nuclei Area
0.0351
G
Regulation of Lysosomal Catabolism in Human Astrocytes for APOE4 vs. APOE3/3
LRP1
LDLR
ATP13A2
log[FC]
H
Regulation of Oxidative Stress-Induced Cell Death in Human Astrocytes for APOE4 vs. APOE3/3 Individuals
HSPB1
NFE2L2
GPR37L1
SOD1
P4HB
ATF4
SFPQ
PARK7
RIPK1
FBXO7
NUP93
PINK1
UBQLN1
LRRK2
SIRT1
NME5
PDE8A
PRKN
log[FC]
I
Consequences of APOE4 Astrocytes on Lysosomes in Brain Niche
APOE3 miBrain
APOE3 miBrain with APOE4 Astrocytes
Lysosome Live Tracker / Astrocytes / Hoechst
J
i
Astrocyte-Lysosome Area Normalized by Astrocyte Area
0.0003
ii
Total-Lysosome Area Normalized by Nuclei Area
0.0003
iii
Total Lysosomes Particle Intensity
0.0007
K
Lipid Droplets in Astrocytes in APOE3 miBrains
APOE3 Astrocytes
APOE4 Astrocytes
BODIPY / GFAP / Hoechst
L
Lipid Droplet Number
<0.0001
APOE3
APOE4
Astrocytes in APOE3 miBrain

**Fig. S15: *APOE4* Astrocyte Dysregulation in miBrain.** (**A**) Gene expression counts from 2D astrocyte monocultures from previously published RNA-seq dataset (*48*) for (i) *GFAP* and (ii) S100β, (**B**) S100b immunoreactivity in miBrains treated with 100 μg/mL LPS for 48 hours compared to untreated miBrains (green: S100b, blue: Hoechst; scale bar, 50 μm), (**C**) quantification of cellular mean intensity of S100b-positive cells in untreated and 100 μg/mL LPS treatment for 48 hours (n = 70 cells/ group; reported as mean and S.E.M., statistical significance via t test), (**D**) GFAP immunoreactivity in miBrains treated with LPS compared to untreated miBrains (green: GFAP, blue: Hoechst; scale bar, 50 μm), (E) quantification of cellular mean intensity of GFAP-positive cells in untreated and LPS-treated conditions (n = 40 cells/ group; reported as mean and standard deviation, statistical significance via t test), (**F**) peroxynitrite (ONOO) in miBrains with (**i**) *APOE3* (left) versus *APOE4* (right) astrocytes incorporated into otherwise-*APOE3* miBrains (green: ONOO, red: live tubulin label, gray: RFP-astrocytes, blue: Hoechst; scale bars, 50 μm), and (**ii-iv**) quantification of peroxynitrite in terms of (**ii**) mean intensity of peroxynitrite in astrocyte-colocalized regions, (**iii**) overall mean intensity of peroxynitrite normalized by nuclei area, and (**iv**) mean intensity of peroxynitrite in tubulin-colocalized region normalized by nuclei area (data from n = 3 replicates and images from n = 9 max projections from confocal z-stacks; statistical analysis via t test; experiment was conducted in 3 independent trials), (**G,H**) DEGs in human astrocytes from snRNAseq dataset (*17*) in *APOE3/4* and *APOE4/4* versus *APOE3/3* individuals (positive log[FC] is upregulated in *APOE4*) for pathway genes in (**G**) regulation of lysosomal protein catabolic process (GO:1905165) and (**H**) regulation (GO:1903201), positive regulation (GO:1903209), and negative regulation (GO:1903202) of oxidative stress-induced cell death, (**I**) lysosomes in miBrains with *APOE3* (left) versus *APOE4* (right) astrocytes incorporated into otherwise-*APOE3* miBrains (green: lysosome live tracker, red: RFP-astrocytes, blue: Hoechst; scale bars, 50 μm), and (**J**) quantification of lysosomes in terms of (**i**) area of lysosomes co-localized with astrocytes normalized by astrocyte area, (**ii**) total segmented lysosome area normalized by nuclei area, and (**iii**) intensity of segmented lysosome "particles" (data from n = 3 replicates and images from n = 9 fields of view; statistical analysis via t test; experiment was conducted in 3 independent trials), (**K**) lipid droplets in *APOE3* versus *APOE4* astrocytes in otherwise *APOE3* miBrains (green: BODIPY, red: GFAP, blue: Hoechst; scale bar, 30 μm), and (**L**) quantification of lipid droplet number for *APOE3* versus *APOE4* astrocytes in *APOE3* miBrains (n = 3 miBrains/ group; results shown on a per-cell basis; statistical significance via t test).

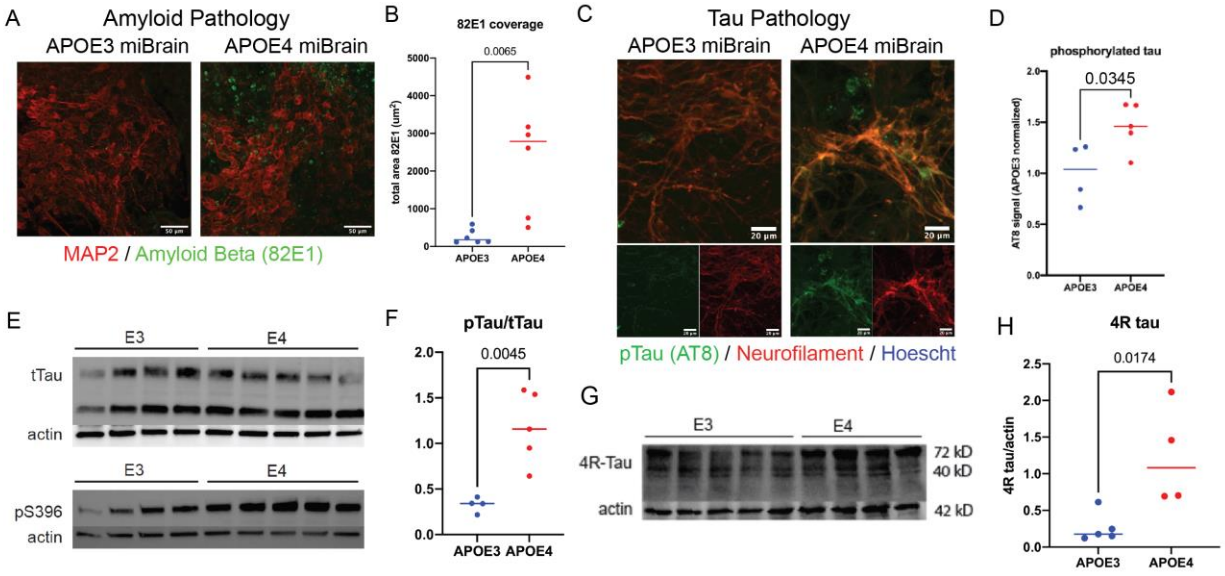


**Fig. S16: Endogenous amyloid and tau pathologies in *APOE4* miBrain.** (**A**) Amyloid aggregate accumulation in *APOE4* (right) compared to *APOE3* (left) miBrains at week 14 (green: amyloid 82E1, red: MAP2; scale bars, 50 µm), (**B**) quantification of total area of 82E1-positive aggregates normalized to cell density in the field of view (n = 6; student's unpaired t test, p = 0.0065), (**C**) immunoreactivity to hyperphosphorylated tau in neurons in *APOE3* (left) versus *APOE4* miBrains (right) at 14 week (green: AT8, red: neurofilament, blue: Hoechst; scale bars, 50 µm), (**D**) quantification of hyperphosphorylated tau as intensity of AT8 signal co-localized with neurofilament-positive cells, normalized to the *APOE3* condition (n = 4; student's t test, p = 0.034), (**E**) western blot analysis for total tau (tTau) and pS396 for *APOE3* and *APOE4* miBrains, using actin as a housekeeping probe for normalization, (**F**) quantification of pS396 between *APOE3* and *APOE4* miBrains, plotted as pTau normalized by tTau, (**G**) western blot analysis for 4R-Tau for *APOE3* and *APOE4* miBrains, using actin as a housekeeping probe for normalization, and (**H**) quantification of 4R-Tau bands.

**Movie S1.**

Visualization of neurovascular units and microglial integration throughout the miBrain with 3D reconstructions (cyan: neurons via tubulin label, red: BMECs, green: iMG via membrane label).

**Movie S2.**

Visualization of astrocytes integrated with neurons and with the blood-brain barrier throughout the miBrain with 3D reconstructions (green: mCherry-astrocytes, cyan: neurons via tubulin label, blue: Hoechst, red: ZO1-BMECs,).

**Movie S3.**

Visualization of oligodendroglia integrated with neurons and Fluoromyelin throughout the miBrain neurovascular units and microglial integration with 3D reconstructions (green: mCherry-oligodendroglia, red: FluoroMyelin, cyan: neurons via tubulin label).
